# The pyruvate branch point controls lymphoid cancer cell dissemination

**DOI:** 10.64898/2026.03.16.712182

**Authors:** Hamidullah Khan, Steven John, Sushmita Roy, Mohd Farhan, Nguyet Minh Hoang, Patrick Buethe, Aman Prasad, Aman Nihal, David T. Yang, Lixin Rui, Jing Fan, Stefan M. Schieke

**Affiliations:** Department of Dermatology, University of Wisconsin-Madison, Madison, WI 53705, USA; MedStar Health Research Institute, Washington, DC 20010, USA; Washington DC VA Medical Center, Washington, DC 20422, USA; Lombardi Comprehensive Cancer Center, Georgetown University, Washington, DC 20057, USA; Morgridge Institute for Research, Madison, Wisconsin, 53706 USA; Cellular and Molecular Biology Graduate Program, University of Wisconsin-Madison, Madison, WI; Department of Medicine, University of Wisconsin School of Medicine and Public Health, Madison, WI, 53792, USA; Novartis Institutes for BioMedical Research, Cambridge, MA 02139; Department of Dermatology, University of Florida, Gainesville, FL 32606 USA; Department of Pathology and Laboratory Medicine, University of Wisconsin-Madison, Madison, WI 53705, USA; Department of Medical Microbiology and Immunology, University of Wisconsin-Madison, 53706; Department of Dermatology, Georgetown University Medical Center, Washington, DC 20057, USA

**Keywords:** lymphoid malignancies, mitochondria, pyruvate, TCA cycle, metabolic reprogramming, redox signaling

## Abstract

Cancer cell dissemination critically determines clinical prognosis, yet metabolic dependencies and corresponding therapeutic targets during spread of lymphoid malignancies remain poorly understood. Here we show that the pyruvate branch point operates as a metabolic checkpoint for lymphoid cancer cell migration and disease dissemination through mitochondrial ROS (mROS)/HIF-1a signaling. Isolation of highly migratory mROS^hi^ cells led us to identify selective metabolic requirements of malignant lymphocyte migration and disease dissemination. Highly migratory cells show a reprogrammed metabolic profile characterized by increased glucose uptake and reduced glucose-carbon entry into the TCA cycle. Reprogramming of the TCA cycle with downregulation of citrate synthase provide the mechanistic basis for decreased pyruvate oxidation leading to increased migration and disease dissemination through mROS/HIF-1a signaling. Our findings connect central carbon metabolism and migratory capacity of lymphoid cancer cells and identify the pyruvate branch point as a metabolic switch and potential therapeutic target in lymphoid cancer cell dissemination.

## INTRODUCTION

Cancer cells exhibit dynamic metabolic profiles that support varying demands during disease progression across different tissue environments.^1–3^ This context-dependent metabolic flexibility presents a major therapeutic challenge for the design of effective metabolic treatments. Identifying limiting requirements and critical branch points in the metabolic network of cancer cells could enable more targeted therapeutic strategies.^4–7^ Pyruvate flux and the balance between glycolysis and oxidative metabolism represent one such branch point, and its regulatory role is well established in various solid cancers as well as in intestinal and hair follicle stem cells.^7–10^ Notably, the effect of pyruvate flux is cancer type-dependent with tumorigenesis in colon cancer relying on increased conversion of pyruvate to lactate and prostate cancer growth being supported by enhanced pyruvate oxidation in the TCA cycle.^11,12^ Downstream of this branch point, several studies have identified TCA cycle reprogramming, specific metabolites, and citrate synthase function as critical regulators in solid cancers.^3,13–15^

Mitochondria are essential organelles that serve as the primary site of cellular energy production while also playing critical roles in biosynthesis and redox signaling across cancer types.^16–19^ Reactive oxygen species (ROS) generated by the mitochondrial electron transport chain (ETC) function as pivotal signaling molecules that regulate gene expression and cellular metabolism, thereby promoting cancer cell proliferation, survival, and metastasis.^20,21^ Cancer cells maintain ROS within a narrow range that supports signaling without causing overt damage, a balance critical for tumor growth and progression.^22,23^ However, studies on mitochondrial metabolism and mROS signaling during solid cancer progression and metastasis have yielded contradictory results, with evidence supporting both metastasis-promoting and metastasis-suppressing effects of ROS, likely depending on factors such as cell or tissue of origin, disease stage, and tissue microenvironment.^24^ In some cancer models, including murine melanoma and pancreatic ductal adenocarcinoma, ROS promote progression and metastasis through activation of Src, Pyk2, or ERK.^25,26^ In other models, such as lung cancer and murine and human melanoma, ROS accumulation can inhibit cell migration and invasion; accordingly, metastatic cells often show upregulation of antioxidant defenses such as the pentose phosphate pathway or decreased mitochondrial ROS production.^23,27–31^

Although a complex picture of metabolic reprogramming and nutrient dependencies has emerged for various solid cancers, the metabolic requirements that support bioenergetic and biosynthetic demands in lymphoid cancers remain less well understood. Normal lymphocytes undergo dynamic metabolic reprogramming to support differentiation, proliferation, and effector functions.^32–34^ These cells utilize diverse nutrients including glucose, lactate, acetate, fatty acids, and glutamine to fuel metabolic pathways that optimize their function in infection, inflammation, and cancer. While glucose can support biosynthetic pathways through the initial steps of glycolysis, fueling of the TCA cycle and oxidative phosphorylation has been shown to be critical for T lymphocyte function and migration, in part mediated through redox signaling and ATP generation.^35–39^ Reactive oxygen species, particularly mitochondrial ROS (mROS), serve as key signaling molecules that modulate lymphocyte function and immune responses.^21,40^ During T cell receptor stimulation, transient increases in mROS promote NFAT activation and IL-2 production, facilitating T cell activation and proliferation.^41^ Conversely, dysregulated ROS production can impair lymphocyte function, promoting exhaustion phenotypes characterized by reduced effector capacity and increased expression of inhibitory receptors.^42^ While growing knowledge of normal lymphocyte metabolism and nutrient requirements is enabling the modulation of immune responses in infection, inflammation, and cancer, deeper insight into the metabolic properties of lymphoid cancer cells would provide a basis for selectively targeting malignant lymphocytes while preserving the host immune response.

In this study, we identify the pyruvate branch point as a critical metabolic checkpoint that regulates lymphoid cancer cell migration and dissemination through mitochondrial signaling. We show that mROS levels support the invasive potential of malignant lymphocytes, with highly invasive cells exhibiting reduced glucose carbon allocation to the TCA cycle. Mechanistically, decreased citrate synthase expression drives this metabolic shift by limiting glucose-derived pyruvate flux into the TCA cycle and elevating mROS/HIF-1a signaling. We establish citrate synthase as a key determinant of invasive capacity, demonstrating that its downregulation promotes lymphoid cancer cell migration through the mROS/HIF-1a axis. These findings reveal how metabolic reprogramming at the pyruvate branch point controls lymphocyte dissemination and identify potential therapeutic targets for limiting the progression of lymphoid malignancies.

## RESULTS

### Mitochondrial ROS promote migration and dissemination of malignant lymphocytes

To investigate the role of mitochondrial signaling and metabolic programming in the migration and *in vivo* dissemination of lymphoid cancer cells, we exploited intrinsic variations of mROS levels to isolate subpopulations with distinct redox signaling profiles. We employed fluorescence-activated cell sorting (FACS) to isolate cells with low and high levels of mROS, mROS^lo^ and mROS^hi^, as detected by the mitochondrial fluorogenic dye MitoSOX Red. To determine whether these subpopulations exhibited phenotypic differences in migration and dissemination capabilities, we subjected them to transwell migration assays and mouse xenograft studies as illustrated in Figure 1A.

**Figure 1.**
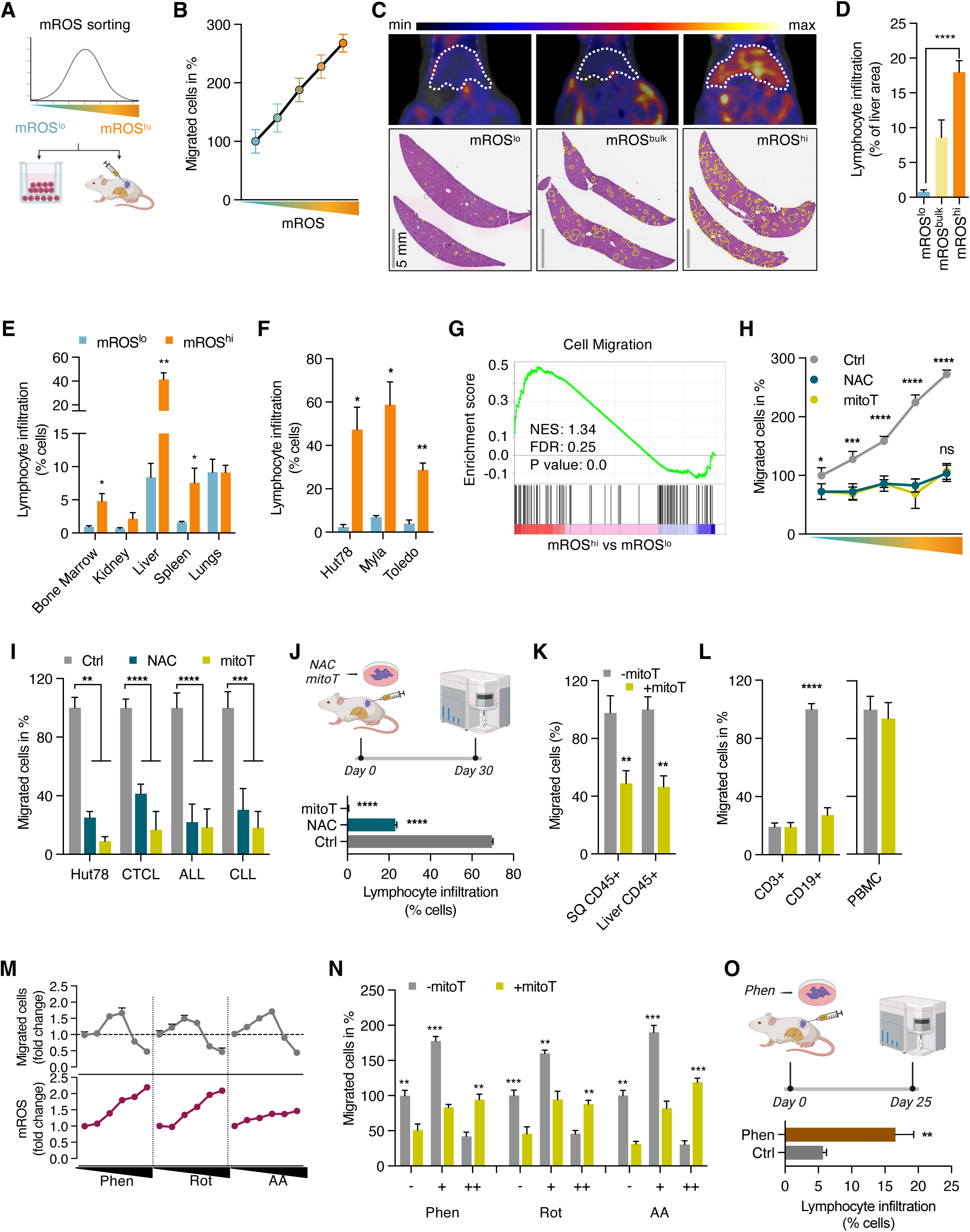
Mitochondrial ROS promote migration and dissemination of malignant lymphocytes (A) Schematic representation of workflow of mROS sorting and subsequent transwell migration and xenograft studies. (B) Transwell migration of Jurkat cells sorted for five different mROS levels ranging from low to high as indicated in different colors. (C) Representative images of ^18^FDG-PET/CT studies and hematoxylin and eosin (H&E)-stained liver sections of hepatic dissemination of sorted mROS^lo^, mROS^bulk^, and mROS^hi^ Hut78 cells. Liver outlined by dotted white lines in PET/CT images and infiltrating lymphocyte aggerates outlined in yellow in H&E images (n=4 mice per group in PET/CT studies). (D) Quantification of hepatic infiltration of sorted Hut78 cells in H&E sections shown as percent of liver section area (n=2 mice). (E) Quantification of organ infiltration after xenografting of HH cells. Hepatic spread was quantified by flow cytometry using anti-hCD45 antibodies (n=4 mice). (F) Hepatic infiltration after second-generation xenografting of isolated mROS^lo^ and mROS^hi^ xenograft tumor cells (n=3 mice/group). (G) Gene set enrichment analysis of FACS-sorted mROS^lo^ and mROS^hi^ Jurkat cells. (H) Effect of antioxidants, N-acetylcysteine (NAC) and mitoTEMPO (mitoT), on migration of Jurkat cells sorted for different mROS levels ranging from low to high. (I) Effect of N-acetylcysteine (NAC) and mitoTEMPO (mitoT) on transwell migration of Hut78 cells and freshly isolated leukemic cells from patients with CTCL (n=4), ALL (n=14), and CLL (n=11). (J) Hepatic dissemination of Hut78 cells pretreated with NAC and mitoT. Hepatic spread was quantified by flow cytometry using anti-hCD45 antibodies (n=3 mice/group). (K) Effect of mitoTEMPO on *ex vivo* migration of xenograft cells isolated from subcutaneous primary tumors (SQ CD45+) and liver infiltrates (Liver CD45+). (L) Effect of antioxidant mitoTEMPO on *ex vivo* migration of freshly isolated CD3+ and CD19+ cells from CLL patients (n=6) and PBMC from healthy subjects (n=10). (M) Levels of mROS and transwell migration of Jurkat cells treated with increasing concentrations of phenformin (0, 1, 2, 4, 6, 10 uM), rotenone (0, 10, 50, 100, 200, 500 nM), and antimycin A (0, 25, 50, 100, 200, 400 nM) for 24 hours. (N) Effect of antioxidant mitoTEMPO on transwell migration of Jurkat cells treated with low (+) and high (++) concentrations of each phenformin (4 and 10 uM), rotenone (50 and 500 nM), and antimycin A (100 and 500 nM). (O) Hepatic dissemination of Hut78 cells pretreated with phenformin. Hepatic spread was quantified by flow cytometry using anti-hCD45 antibodies (n=4 mice/group). All data presented as mean ± SD of at least triplicate measurements; ns, non-significant. ∗p < 0.05, ∗∗p < 0.01, ∗∗∗p < .001, ∗∗∗∗p < 0.0001.

First, we assessed the migration potential of the isolated subpopulations in matrigel-coated transwell assays. We observed a striking correlation between cellular mROS levels and migration potential within the isolated subpopulations (Figure 1B). Cells with the highest level of mROS, mROS^hi^, showed a markedly enhanced migration potential (eMP) compared to the four other isolated subpopulations with lower levels of mROS. Subsequently, we focused on mROS^lo^ and mROS^hi^ cells to delineate the role of redox signaling and metabolic requirements underlying migration of malignant lymphocytes. Notably, mROS^lo/hi^ states were not stable but dynamic in nature. After 24 hours in culture post-sorting, the isolated subpopulations exhibited similar mROS levels (Figure S1A) and migration potential (Figure S1B), indicating the dynamic nature of mROS generation similar to previously described variations in mitochondrial membrane potential.^43,44^

Next, we tested whether the increased migratory properties of mROS^hi^ cells observed in transwell assays correlated with increased dissemination and solid organ infiltration *in vivo*. Transplantation of mROS^hi^ cells led to markedly increased liver infiltration compared to both mROS^lo^ and bulk cells. ^18^FDG-PET/CT scans revealed increased hepatic FDG uptake in animals receiving mROS^hi^ cells compared to those receiving mROS^lo^ and sham-sorted bulk cells, mROS^bulk^ (Figure 1C). This finding was confirmed by H&E staining of liver sections demonstrating increased hepatic lymphocyte infiltration of xenografted mROS^hi^ cells compared to mROS^lo^ and bulk cells (Figures 1C, 1D, S1C). Notably, we also observed increased dissemination of the mROS^hi^ cells into bone marrow, spleen, and kidneys indicating that increased dissemination is not limited to the hepatic microenvironment (Figure 1E). Moving forward, we chose to focus on hepatic spread given that it was most pronounced in our xenograft model and is a common feature of lymphoid malignancies.^45,46^ Despite marked differences in mROS levels, growth of primary tumors initiated from mROS^lo^, mROS^hi^, and bulk subpopulations did not show any significant differences (Figure S1D).

To confirm that the enhanced migratory mROS^hi^ phenotype also exists in mouse xenograft tumors, we isolated mROS^lo^ and mROS^hi^ cells from established xenograft tumors. Similar to cells isolated from cell culture, mROS^hi^ xenograft tumor cells demonstrated increased migration *ex vivo* (Figure S1E) and hepatic infiltration after serial transplantation (Figure 1F). At the same time, we observed no differences in the growth of second-generation primary tumors (Figure S1F).

These data demonstrate that isolated mROS^hi^ cells show enhanced migration and dissemination across various tissues *in vivo*. Consistent with this more aggressive phenotype, the mROS^hi^ cell subpopulation exhibited an enriched transcriptional signature for genes involved in cell migration (Figure 1G).

Next, we elucidated the functional relationship between mROS levels and migration in malignant lymphocytes. Treatment with pan-ROS scavenger N-acetylcysteine (NAC) and the mitochondrial-targeted antioxidant mitoTEMPO abolished the migratory advantage of isolated mROS^hi^ cells suggesting a critical role of mROS supporting the migratory phenotype (Figure 1H). Notably, antioxidants reduced migration to a similar level in all isolated subpopulations suggesting that increased migration observed with rising levels of mROS is primarily driven by redox signaling. We confirmed the role of mROS for malignant lymphocyte migration by using another fluorogenic probe, CellROX Green, to isolate mROS^lo/hi^ cells which yielded similar results effectively ruling out any dye-specific artifacts (Figure S2A). Consistent with the anti-migratory effect of antioxidants in sorted mROS subpopulations, we found a marked anti-migratory effect of NAC and mitoTEMPO in unsorted cell lines and freshly isolated leukemic cells from patients with Sézary syndrome, a leukemic form of cutaneous T cell lymphoma (CTCL), B and T acute lymphoblastic leukemia (ALL), and chronic lymphocytic leukemia (CLL), indicating the universal role of mROS in malignant lymphocyte migration across a spectrum of various lymphoid malignancies (Figure 1I). We further validated the anti-migratory effect of antioxidants in several additional B- and T-lymphoid cancer cell lines (Figure S2B). Importantly, the anti-migratory effect of antioxidants was not due to suppression of cell proliferation and viability (Figure S2C).

Next, we asked whether antioxidants lead to decreased hepatic infiltration *in vivo*. Transplantation of cells treated with NAC and mitoTEMPO for 24 hours prior to injection led to markedly diminished hepatic infiltration (Figure 1J) without effects on primary tumor growth (Figure S2D). Similar results were obtained by treatment of mice with NAC or mitoTEMPO confirming the anti-migratory effect of antioxidant treatment *in vivo* (Figures S2E and S2F).

Given the strong relationship between mROS and malignant lymphocyte dissemination, we asked whether the pro-migratory role of mROS is maintained during disease spread and carried over to new tumor sites and microenvironments rather than being unique to the primary tumor injection site. To examine this, we isolated xenograft cells from subcutaneous primary tumors and hepatic infiltrates out of the same animals. In *ex vivo* transwell assays, we could show that the pro-migratory effect of mROS was retained in tumor cells that had spread to the liver (Figure 1K) indicating its persistence in various tissue microenvironments.

To test whether this feature is unique to malignant lymphocytes or also applies to healthy lymphocytes, we isolated leukemic cells expressing CD19 and non-leukemic CD3+ T lymphocytes from CLL patients. We examined these subpopulations independently and found that only malignant lymphocytes exhibited decreased migration in response to mitoTEMPO, indicating a selective role of mROS in malignant lymphocyte migration (Figure 1L). Similarly, mitoTEMPO had no effect on transwell migration of isolated peripheral blood mononuclear cells (PBMC) from healthy donors (Figure 1L). These data suggests that mROS is a critical driver of migration and dissemination selective for malignant lymphocytes.

Next, we examined whether an increase in mROS through modulation of either complex I or complex III of the mitochondrial electron transport chain changes the migratory phenotype. Complex I inhibitors, phenformin and rotenone, and complex III inhibitor, antimycin A, increased levels of mROS in a dose-dependent fashion (Figure 1M, lower panels). This led to an increase in migration at modest elevations of mROS with low concentrations of ETC inhibitors followed by a decline in migration at further increased levels of mROS with higher concentrations of ETC inhibitors (Figure 1M, upper panels). Scavenging ROS with mitoTEMPO led to decreased migration at modestly increased mROS levels while restoring migration to baseline at higher mROS levels (Figures 1N and S2G). This suggests a biphasic, concentration-dependent role of mROS with a pro-migratory effect at modestly elevated levels and higher levels showing a suppressive effect on migration. Both concentrations of ETC inhibitors showed no changes of cellular ATP levels ruling out an energetic deficiency at higher, migration-suppressive concentrations (Figure S2H). Consistent with the pro-migratory effect of mitochondrial complex I inhibition, cells pretreated with phenformin demonstrated increased hepatic infiltration in xenograft studies (Figure 1O).

These findings indicate that migration and *in vivo* dissemination of malignant lymphocytes are driven by changes in mROS levels. This allows for the isolation of cells with enhanced migratory potential (eMP-mROS^hi^ cells) to delineate metabolic and signaling mechanisms underlying malignant lymphocyte migration and spread.

### Pro-migratory effect of mROS is mediated through HIF-1a

Given that mROS is a well-known activator of HIF-1a during hypoxia or mitochondrial complex I inhibition by biguanides, we hypothesized that the pro-migratory effect of mROS is mediated by HIF-1a.^5^ First, we showed that eMP-mROS^hi^ cells isolated from various cell lines and primary leukemic samples display increased levels of HIF-1a (Figures 2A-C). Furthermore, we were able to demonstrate a robust correlation between mROS and HIF-1a levels highlighting the potential role of the mROS/HIF-1a axis in the regulation of malignant lymphocyte migration (Figure S3A).

**Figure 2.**
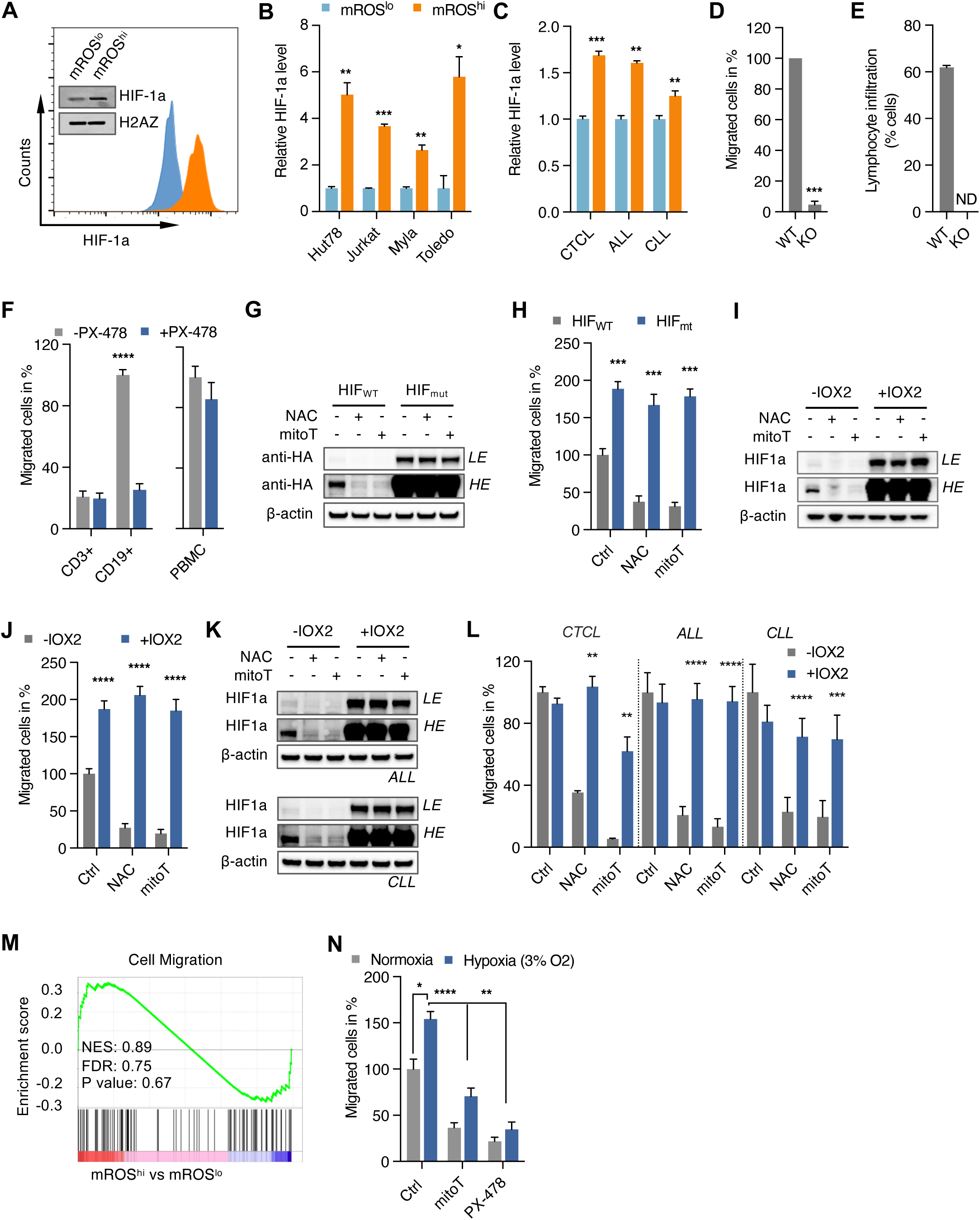
Promotion of malignant lymphocyte spread by mROS is mediated through HIF-1a (A) Levels of HIF-1a in sorted mROS^lo^ and mROS^hi^ Hut78 cells detected by flow cytometry and immunoblotting. Immunoblots from nuclear extracts with antibody against H2A-Z used as loading control. (B) Flow-cytometric quantification of HIF-1a in mROS^lo^ and mROS^hi^ cells of indicated cell lines. HIF-1a levels shown in mean fluorescence intensity. (C) Increased HIF-1a levels in isolated leukemic cells from patients with CTCL (n=2), ALL (n=3), and CLL (n=3). (D and E) Effect of HIF-1a knockout in Hut78 cells on invasive potential in transwell assays (D) and hepatic infiltration *in vivo* (E; n=3 animals per group). (F) Effect of PX-478 on *ex vivo* migration of freshly isolated CD3+ and CD19+ cells from CLL patients (n=6) and PBMC from healthy subjects (n=10). (G and H) Immunoblots (G) and corresponding transwell migration assay (H) of Jurkat cells transfected with either wild-type HA-tagged HIF-1a (HIF_WT_) or mutant HA-tagged-HIF-1a P402A/P564A (HIF_mut_) followed by treatment with antioxidants, NAC or mitoTEMPO. LE, low exposure, HE, high exposure. (I and J) Immunoblots (I) and corresponding transwell migration assays (J) of Jurkat cells treated with PHD inhibitor, IOX2, and either NAC or mitoTEMPO. LE, low exposure, HE, high exposure. (K and L) Immunoblots (K) and corresponding e*x vivo* transwell migration assays (L) of leukemic cells treated with IOX2 and either NAC or mitoTEMPO after isolation from patients with CTCL (n=3), ALL (n=8), CLL (n=8). LE, low exposure, HE, high exposure. (M) Gene set enrichment analysis of Jurkat cells treated with PX-478 and sorted into mROS^lo^ and mROS^hi^ subpopulations. (N) Effect of hypoxia (3% O_2_) versus normoxia on transwell migration of Jurkat cells treated with mitoTEMPO or PX-478. All data presented as mean ± SD of at least triplicate measurements; ns, non-significant. ∗p < 0.05, ∗∗p < 0.01, ∗∗∗p < .001, ∗∗∗∗p < 0.0001.

Using HIF-1a knockout (KO) lines generated with the CRISPR/Cas9 system, we showed that HIF-1a deletion markedly reduced transwell migration (Figure 2D) and hepatic infiltration *in vivo* (Figure 2E) indicating that HIF-1a is necessary for migration and dissemination of malignant lymphocytes. We confirmed the pro-migratory effect of HIF-1a in various malignant B and T lymphoid cells lines using the small-molecule HIF-1a inhibitor, PX-478 (Figures S3B and S3C). Consistent with the selective role of mROS in malignant lymphocyte migration, we found that the reduced migration in response to HIF-1a suppression is selective for malignant cells sparing normal T cells from CLL patients (Figure 2F). Similarly, we did not observe an effect of HIF-1a suppression on migration of isolated PBMCs from healthy donors (Figure 2F) further underlining the selective role of mROS and HIF-1a in the migration of malignant lymphocytes. Importantly, neither genetic nor pharmacologic suppression of HIF-1a decreased mROS levels excluding an indirect effect through reduction of mROS (Figures S3D and E).

These data suggest that HIF-1a is downstream of mROS in regulating migration of malignant lymphocytes. To explore this, we tested whether HIF-1a activation can restore migration in antioxidant-treated cells. Expression of prolyl hydroxylation-deficient HIF-1a (Figures 2G and H) and pharmacologic activation through the prolyl hydroxylase inhibitor IOX-2 (Figures 2I and J) effectively restored HIF-1a levels and migration in cells with disrupted mROS signaling. We validated these findings in isolated leukemic cells from patients with CTCL, B- and T-ALL, and CLL (Figures 2K and L). These results support the hypothesis that HIF-1a is a critical downstream mediator of mROS in the regulation of the migratory phenotype. Consistent with this, mitoTEMPO treatment during HIF-1a suppression did not further lower migration (Figure S3F). A GSEA analysis of mROS^hi^ vs. mROS^lo^ cells demonstrated that treatment with the small-molecule HIF-1a inhibitor, PX-478, abrogated the pro-migratory transcriptional profile of the eMP cells (Figure 2M). Consistent with the role of the mROS/HIF-1a axis in cell migration, activation of HIF-1a through hypoxia also led to increased migration which was antioxidant- and PX-478-sensitive (Figure 2N).

In summary, our data indicate that the mROS/HIF-1a axis is a critical regulator of malignant lymphocyte dissemination. Given that ETC activity and mROS generation are embedded into a complex metabolic network, we next asked which fuel choices and metabolic programming influence the migratory behavior of lymphoid cancer cells through modulating mROS/HIF-1a activity.

### Glucose is an essential fuel for migration through activation of mROS/HIF-1a

To elucidate fuel preferences and metabolic requirements of malignant lymphocyte migration, we performed untargeted metabolomic analyses which showed broad differences in metabolic programming between eMP-mROS^hi^ and mROS^lo^ cells (Figure 3A). More specifically, enhanced migratory cells showed increased levels of various glycolytic intermediates (Figure S4A). Compared to mROS^lo^ cells, isolated eMP-mROS^hi^ cells showed more pronounced differences in the extracellular acidification rate (ECAR) than in the oxygen consumption rate (OCR) suggestive of an enhanced glycolytic phenotype (Figures 3B, S4B and C). Moreover, glucose consumption was markedly increased in eMP-mROS^hi^ cells across several different lymphoid cancer cell lines in xenograft tumors, cell culture, and isolated leukemic lymphocytes (Figures 3C and D, S4D).

**Figure 3.**
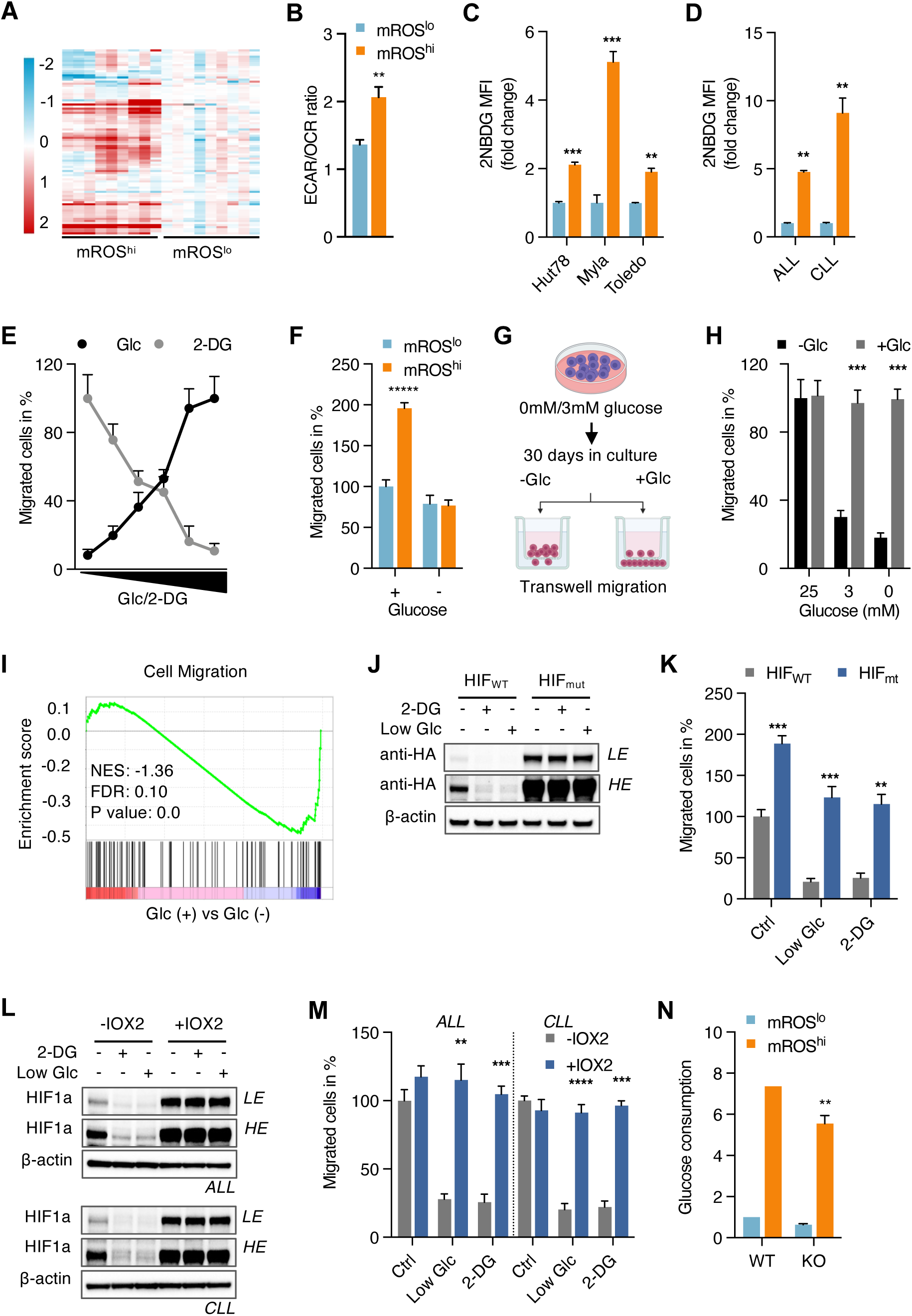
Glucose is an essential fuel for migration through activation of mROS/HIF-1a (A) Clustering heatmap of metabolomic profiles of sorted mROS^lo^ and mROS^hi^ Jurkat cells (n=3 independent samples analyzed in triplicates). (B) Ratio of mitochondrial oxygen consumption rate (OCR) and extracellular acidification rate (ECAR) in isolated mROS^lo^ and mROS^hi^ cells. (C and D) Glucose uptake observed in mROS^lo^ and mROS^hi^ cells isolated from mouse xenograft tumors (C; n=3 tumors of each cell line) and isolated leukemic cells (D; ALL, n=3 patients; CLL, n=3 patients). (E) Effect of glucose-limiting conditions or inhibition of glycolysis with 2-deoxyglucose (2-DG) on transwell migration. Cells were incubated with different concentrations of glucose or 2-DG (0, 2.5, 5, 10, 20, 25 mM for both) during migration in transwell chambers. (F) Effect of glucose-limiting conditions on migration of isolated mROS^lo^ and mROS^hi^ cells. (G and H) Effect of long-term culture in glucose-limiting media on transwell migration (G). Transwell assays were performed after 30 days in glucose-limiting conditions with unchanged glucose concentration (-Glc) or freshly added glucose during the 24 h migration studies (+Glc; H) (I) Gene set enrichment analysis of Jurkat cells in glucose-free conditions. (J and K) Representative immunoblots (J) and corresponding transwell migration assay (K) of Jurkat cells transfected with either wild-type HA-tagged HIF-1a (HIF_WT_) or mutant HA-tagged-HIF-1a P402A/P564A (HIF_mut_) in media containing 3 mM of glucose or 25 mM of 2-DG. (L and M) Representative immunoblots (L) and corresponding *ex vivo* transwell migration assays (M) of isolated leukemic cells treated with PHD inhibitor IOX2 and NAC or mitoTEMPO (ALL, n=4 patients; CLL, n=4 patients). LE, low exposure, HE, high exposure. (N) Glucose uptake observed in control and HIF1a-KO Hut78 cells. mROS^lo^ in light blue bars; mROS^hi^ in orange bars in (B-D, F, N). All data presented as mean ± SD of at least triplicate measurements; ns, non-significant. ∗p < 0.05, ∗∗p < 0.01, ∗∗∗p < .001, ∗∗∗∗p < 0.0001.

Given that eMP-mROS^hi^ cells show increased glucose uptake, we hypothesized that glucose is a critical fuel for malignant lymphocyte migration through activation of the mROS/HIF-1a axis. To test this hypothesis, we tested the effect of limiting glucose conditions and inhibition of glycolysis on migration. Gradual lowering of media glucose concentration or increasing amounts of 2-deoxyglucose inhibited migration in transwell assays indicating that glucose and glycolytic activity are necessary for malignant lymphocyte migration (Figure 3E). Importantly, we did observe a markedly different migration potential within a glucose range from 2.5 mM (45 mg/dL) to 10 mM (180 mg/dL) which is close to the physiologic range in human plasma. This suggests that fasting and postprandial blood glucose variations could potentially modulate the migratory behavior of malignant lymphocytes. Consistent with glucose as a critical driver of malignant lymphocyte migration, the migratory advantage of eMP-mROS^hi^ cells was abolished in low-glucose conditions (Figure 3F). Next, we tested whether prolonged glucose-limiting conditions would overcome the glucose-dependency of migrating cells (Figure 3G). Interestingly, cells could not adapt their migratory metabolic requirements during prolonged times of glucose-limiting conditions (Figure 3H). Migratory ability was, however, readily restored upon addition of glucose to the culture media (Figure 3H). The inability to adjust metabolic fuel requirements of the migratory state under prolonged low-glucose conditions indicates the critical need of glucose for migration. GSEA analysis confirmed the anti-migratory effect of limiting glucose conditions leading to decreased expression of cell migration genes (Figure 3I).

Based on our observation of the mROS/HIF-1a axis as a critical regulator of migration, we hypothesized that glucose and glycolytic activity drive migration through mROS/HIF-1a signaling. Consistent with this, we observed that low-glucose culture conditions and inhibition of glycolysis led to reduced mROS levels and increased mitochondrial respiration (Figures S4E and F). Overexpression of mutant prolyl hydroxylation-deficient HIF-1a restored HIF-1a levels and migration in low-glucose conditions and during inhibition of glycolysis with 2DG (Figures 3J&K). We observed the same effects with pharmacologic activation of HIF-1a through IOX-2 in low-glucose conditions and presence of 2DG (Figures S4G and H). Those results were confirmed in freshly isolated ALL and CLL cells suggesting that the pro-migratory effect of glucose in malignant lymphocytes is mediated by HIF-1a (Figures 3L and M). Consistent with HIF-1a as a downstream mediator of the effect of glucose and glycolysis on migration, we found the increased glucose uptake of the eMP-mROS^hi^ state to be retained in HIF-1a-knockout cells (Figure 3N). This indicates that the metabolic phenotype with increased glucose uptake of eMP-mROS^hi^ cells is independent of HIF-1a signaling.

Taken together, these findings demonstrate that glucose is an essential fuel for malignant lymphocyte migration through activation of the mROS/HIF-1a axis. Next, we wanted to elucidate critical metabolic nodes in malignant lymphocyte migration and dissemination.

### Migration is supported by TCA cycle changes and altered glucose metabolization

Given the broad differences in metabolic programming and more specifically glucose consumption, we used [U-^13^C]glucose tracing to determine the fate of glucose in eMP-mROS^hi^ cells. Labeled glucose entered cells and the glycolytic pathway almost completely as indicated by similar labeling of approximately 99% of the glucose-6-phosphate pools in both subpopulations (Figure 4A). In contrast, eMP-mROS^hi^ cells showed reduced M+2 labeling of TCA cycle intermediates, including α-ketoglutarate, fumarate, and malate, indicating suppressed glucose-carbon entry via the pyruvate dehydrogenase (PDH) pathway (Figure 4A). This was further supported by a decreased citrate M+2 to pyruvate M+3 ratio in eMP-mROS^hi^ cells (Figure S5A). Concordantly, M+2 labeling of TCA cycle-derived amino acids, aspartate (generated from oxaloacetate) and glutamate (generated from α-KG), was also reduced, consistent with diminished glucose contribution through TCA cycle intermediates. Moreover, suppression of total isotopologue labeling [1-(M+0)] across TCA cycle intermediates indicates decreased overall glucose-carbon contribution including non-PDH pathways irrespective of the number of TCA cycle turns (Figure S5B). Interestingly, eMP-mROS^hi^ cells did not show increased shunting of glucose into the pentose phosphate pathway as detected by 6-phosphogluconate and ribulose-5-phosphate (Figure 4A). We also did not observe a significant difference in NADP+/NADPH and GSH/GSSG ratios suggesting that increased mROS levels in mROS^hi^ cells are not sufficient to cause oxidative stress and a concomitant antioxidant response (Figures S6A and B). This is further supported by the finding that several antioxidant enzymes were not differently expressed in mROS^lo^ and mROS^hi^ cells (Figure S6C).

**Figure 4.**
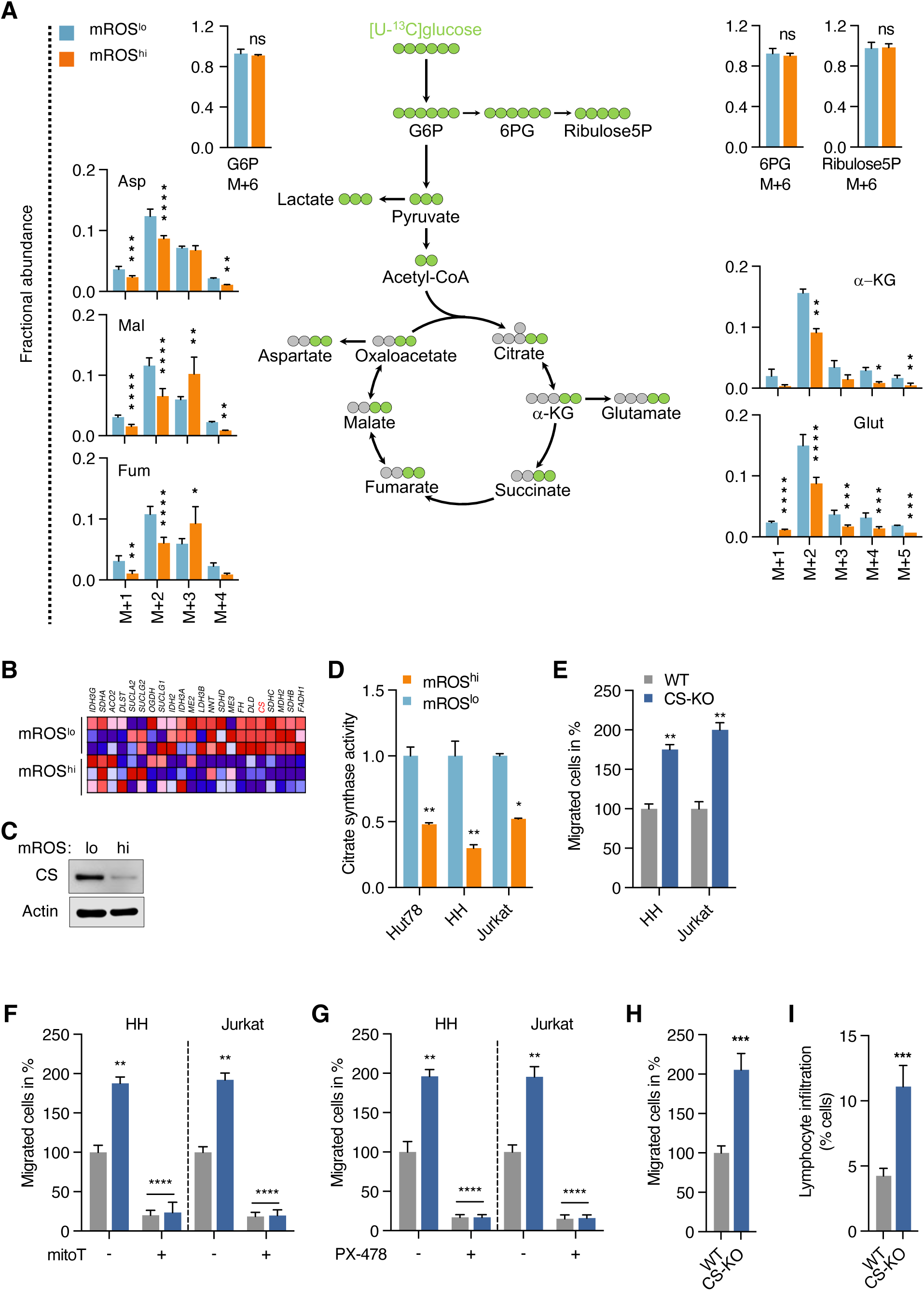
The enhanced migratory phenotype shows TCA cycle changes and altered glucose metabolization (A) Schematic representation of glucose metabolism and isotopologue label distribution from [U-^13^C]glucose in isolated mROS^lo^ and mROS^hi^ Jurkat cells. (B) Clustered heatmap analysis of TCA cycle gene profiles of isolated mROS^lo^ and mROS^hi^ Jurkat cells (performed in technical triplicates). (C) Immunoblots of sorted mROS^lo^ and mROS^hi^ cells probed against citrate synthase with actin levels serving as loading control. (D) Citrate synthase activity in isolated mROS^lo^ and mROS^hi^ cells. (E) Transwell migration assay of wildtype (WT) and citrate synthase knockout (CS-KO) HH cells. (F and G) Effect of mitoTEMPO (F) and PX-478 (G) on transwell migration of WT and CS-KO cells. (H) *Ex vivo* migration of xenograft tumor cells isolated primary tumors grown from WT and CS-KO cells. (I) Hepatic infiltration of WT and CS-KO cells. Wildtype (WT) in grey bars; citrate synthase knockout (CS-KO) in blue bars in (E-I). All data presented as mean ± SD of at least triplicate measurements; ns, non-significant. ∗p < 0.05, ∗∗p < 0.01, ∗∗∗p < .001, ∗∗∗∗p < 0.0001.

These results indicate that metabolic profile and the fate of glucose are intricately linked to the migratory potential of malignant lymphocytes, with eMP-mROS^hi^ cells showing a reduced oxidative phenotype. We next sought to elucidate the molecular basis for reduced shunting of glucose into the TCA cycle in those cells. RNA sequencing analysis revealed broad changes at the level of the TCA cycle with diminished expression of several TCA cycle enzyme genes (Figure 4B). Given its central position at the entry point of acetyl-CoA into the TCA cycle, we focused on citrate synthase as a potential molecular switch in the control of the migratory phenotype.

We found that citrate synthase protein levels (Figure 4C) and enzymatic activity are reduced in eMP-mROS^hi^ cells (Figure 4D). Deletion of citrate synthase (CS-KO) using CRISPR/Cas9 technology (Figure S7A) resulted in increased lactate production and a decreased citrate/pyruvate ratio demonstrating a shift from pyruvate oxidation in the TCA cycle to increased aerobic glycolysis (Figures S7B and C). Notably, in CS-KO cells, we observed increased mROS and HIF-1a levels (Figures S7D and E) along with an enhanced migratory phenotype (Figure 4E) resembling eMP-mROS^hi^ cells.

Enhanced migration in CS-KO cells was sensitive to mitoTEMPO and PX-478 indicating that downregulation of citrate synthase induces migration through mROS/HIF-1a similar to eMP-mROS^hi^ cells (Figures 4F and G). Of note, the glycolytic phenotype as measured by lactate production was preserved in the presence of mitoTEMPO and PX-478 (Figures S7F and G). This indicates that metabolic reprogramming of the pyruvate branch point precedes and drives mROS/HIF-1a signaling, not the reverse. We confirmed the increased migratory phenotype of CS-KO cells isolated from primary tumors *ex vivo* along with increased hepatic infiltration in mouse xenograft studies (Figures 4H and I). At the same time, there was no effect of CS deletion on primary tumor growth as compared to wild-type cells indicating the divergent metabolic needs of primary tumor growth and disease dissemination (Figure S7H).

These data demonstrate that enhanced migratory cells show reduced glucose-carbon contribution to the TCA cycle due to modulated expression of several TCA cycle enzymes. Among those, citrate synthase is a critical regulator of the migration potential of malignant lymphocytes through modulating mROS/HIF-1a. Our data shows that metabolic fuel selection and glucose partitioning are key determinants of the migratory potential of lymphoid cancer cells. Based on our findings, we hypothesized that (i) pyruvate as a downstream metabolite of glucose and (ii) the pyruvate branch point as a metabolic node are critical regulators of malignant lymphocyte migration and *in vivo* dissemination.

### Pyruvate branch point as a metabolic node controlling migratory potential in malignant lymphocytes

Consistent with these hypotheses, supplementation of pyruvate restored migration in glucose-free media in a dose-dependent manner suggesting that pyruvate is indeed a critical metabolite of glucose sufficient to drive migration (Figure 5A). Similar to glucose-containing media, we found that pyruvate supplementation led to activation of mROS/HIF-1a signaling (Figure 5B and C) which was necessary for pyruvate to restore migration in the absence of glucose (Figure 5D and E). We confirmed the pro-migratory effect of pyruvate in glucose-limiting conditions in isolated CLL cells in contrast to normal T cells from the same patients or PBMCs isolated from healthy donors (Figure 5F). These findings demonstrate that pyruvate is a critical metabolite in the glucose-dependent migration of malignant lymphocytes.

**Figure 5.**
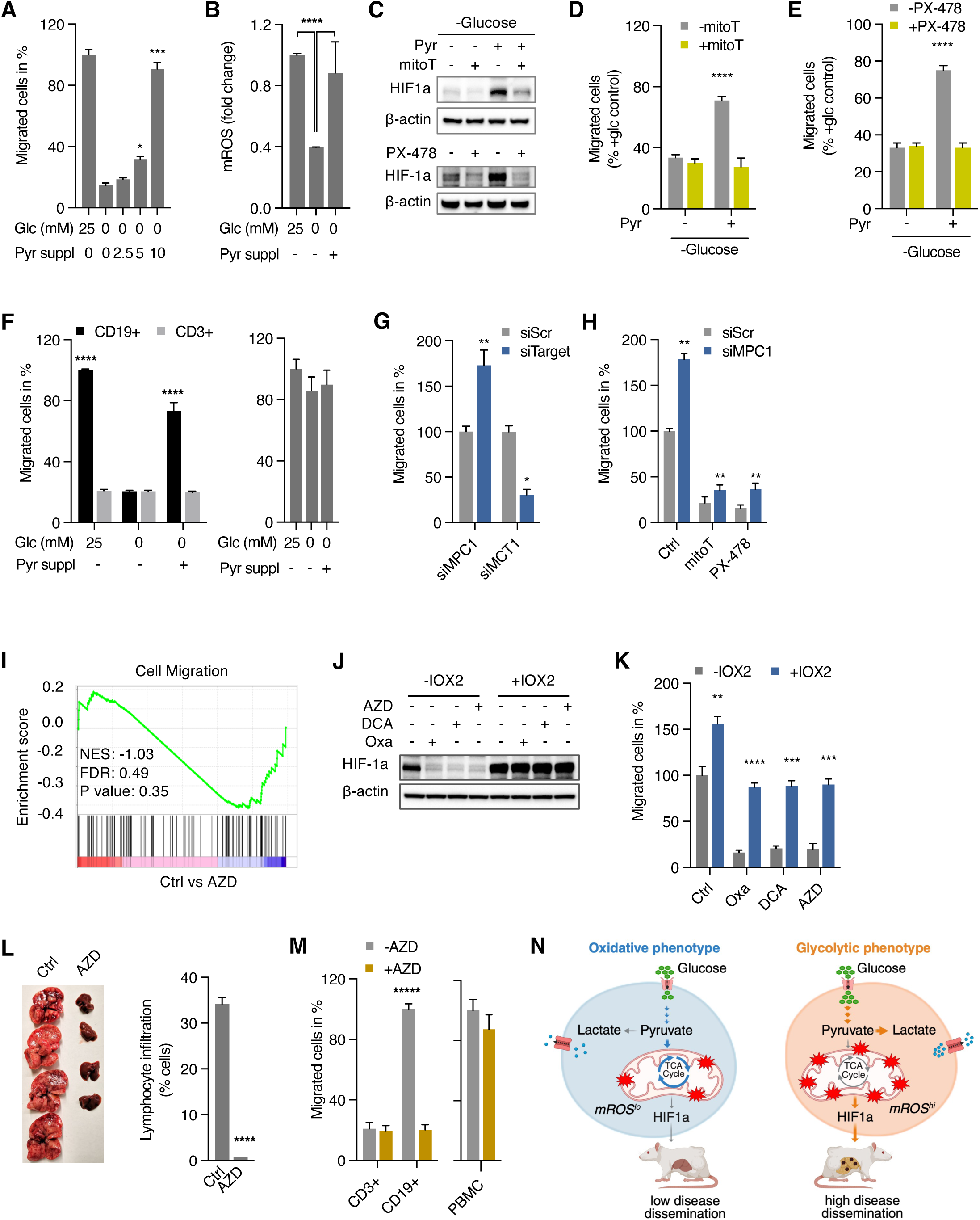
Metabolic control of migratory potential at the pyruvate branch point (A) Transwell migration of cells supplemented with 2.5, 5, 10 mM pyruvate in glucose-free media for 24 hours. (B) Levels of mROS in response to pyruvate supplementation (10mM) in glucose-free media. (C) Immunoblots of HIF-1a in Jurkat cells after pyruvate supplementation in glucose-free media. (D and E) Effect of mitoTEMPO (D) and PX-478 (E) on transwell migration in glucose-free media with and without supplemented pyruvate. (F) Effect of pyruvate supplementation on *ex vivo* migration of freshly isolated CD3+ and CD19+ cells from CLL patients (n=6) and PMBCs from healthy subjects (n=10) in glucose-free media. (G) Transwell migration assay of cells transfected with mitochondrial pyruvate carrier 1 siRNA (siMPC1), monocarboxylate transporter 1 siRNA (siMCT1) or scrambled siRNA (siSCR). (H) Effect of mitoTEMPO (mitoT) and PX-478 on migration of cells transfected with mitochondrial pyruvate carrier 1 siRNA (siMPC1) or scrambled siRNA (siSCR). (I) Gene set enrichment analysis of Jurkat cells treated with AZD3965 compared to untreated control. (J and K) Immunoblots (J) and corresponding transwell migration assays (K) of Jurkat cells treated with PHD inhibitor IOX2 and Oxamate, DCA, or AZD3965. (L) Representative images of livers of mice treated with AZD3965 versus control and corresponding quantitation of hepatic disease infiltration in xenograft studies of Hut78 cells (n=4 animals per group). (M) Effect of AZD3965 on *ex vivo* migration of freshly isolated CD3+ and CD19+ cells from CLL patients (n=6) and PBMC from healthy subjects (n=13). (N) Schematic summary of oxidative versus glycolytic metabolic programming controlling dissemination potential of lymphoid cancer cells through redox signaling. All data presented as mean ± SD of at least triplicate measurements; ns, non-significant. ∗p < 0.05, ∗∗p < 0.01, ∗∗∗p < .001, ∗∗∗∗p < 0.0001.

Next, we sought to explore the role of the pyruvate branch point as a metabolic regulator of migration and xenograft spread. First, we tested the effect of changing the flux of pyruvate through knockdown of the mitochondrial pyruvate carrier (MPC1) or monocarboxylate transporter (MCT1) with decreased or increased pyruvate oxidation, respectively, as measured by lactate production (Figures S8A and B). Shifting pyruvate flux away from the mitochondria through suppression of MPC1 increased migration while increased oxidation of pyruvate in MCT1-knockdown cells decreased migration (Figure 5G).

Consistent with our observation of mROS/HIF-1a signaling mediating the pro-migratory effect of glucose or pyruvate, increased migration of MPC1-knockdown cells was sensitive to mitoTEMPO and PX-478 placing the pyruvate checkpoint upstream of mROS/HIF-1 (Figure 5H). Additionally, pyruvate supplementation of MCT1-knockdown cells did not restore migration ruling out a direct effect of pyruvate on migration and providing further evidence for the critical role of the pyruvate branch point in malignant lymphocyte migration (Figure S8C).

Similarly, pharmacologically increased pyruvate oxidation using oxamate (LDH inhibitor), dichloroacetic acid (DCA; pyruvate dehydrogenase kinase inhibitor activating PDH), and AZD3965 (MCT1 inhibitor) (Figure S8D and E) led to a marked decrease in transwell migration (Figure S8F) suggesting that the fate of pyruvate determines the migratory potential with increased pyruvate oxidation having an anti-migratory effect. The critical role of pyruvate flux in migration was further supported by GSEA analysis demonstrating downregulation of pathways related to cell migration in response to AZD3965 (Figure 5I). Shifting pyruvate flux into the TCA cycle blocked the pro-migratory effect of pyruvate under glucose-free conditions ruling out a branch point-independent effect of pyruvate (Figure S8G). Consistent with the notion that HIF-1a is regulated by the flux of pyruvate, oxamate, DCA, and AZD lowered HIF-1a levels whereas IOX2 restored HIF levels (Figure 5J) and migration during pharmacologically enhanced pyruvate oxidation (Figure 5K).

To examine the effect of increased oxidation of pyruvate in the TCA cycle *in vivo*, we used AZD3965, currently undergoing clinical trials in advanced solid cancer and lymphoma.^47^ Treatment of mice with AZD resulted in a small reduction in primary tumor size (Figure S8H) along with a marked reduction in hepatic dissemination (Figure 5L). Notably, lactate levels and citrate/pyruvate ratio were increased in tumor cell lysates confirming the metabolic, pro-oxidative effect of AZD on tumor cells *in vivo* (Figures S8I and J). We confirmed the anti-migratory effect of increased glucose oxidation in CLL samples. Isolated malignant CD19+ B cells showed reduced migration while normal CD3+ T cells from CLL patients and PBMC isolated from healthy volunteers showed no effect of AZD3965 treatment (Figure 5M).

In summary, our data suggest that pyruvate as a downstream metabolite of glucose and its metabolic fate of oxidation in the TCA cycle or reduction to lactate are critical regulators of malignant lymphocyte migration and *in vivo* dissemination through mROS/HIF-1a signaling. Based on this, we propose that the pyruvate branch point represents a metabolic checkpoint in the process of lymphoid cancer cell migration and dissemination (Figure 5N).

## DISCUSSION

Our study reveals that the pyruvate branch point functions as a metabolic switch for lymphoid cancer cell migration. Reduced pyruvate oxidation in the TCA cycle activates mROS/HIF-1a signaling and a pro-migratory transcriptional program that controls migration of malignant lymphocytes. These findings further define the relationship between metabolism and migration: rather than merely fueling a separately regulated motility program, metabolic reprogramming at the pyruvate branch point actively instructs and controls the migratory phenotype of malignant lymphocytes. Our results demonstrate a pro-migratory role of mROS/HIF-1a in lymphoid cancers which is remarkably consistent across leukemic cell lines and freshly isolated human leukemic CTCL cells, B- and T-ALL, and CLL cells.

Whether mROS promote or suppress cancer cell metastasis has been debated across tumor types.^24^ We observed a biphasic migration response to mROS levels with a pro-migratory effect of moderately elevated levels versus an anti-migratory effect at higher levels of mROS. This biphasic effect might partially explain some of the seemingly contradictory findings of ROS on cancer spread with the net effect of ROS influenced by where a given cell sits along a dose-response curve rather than a simple pro- or anti-metastatic dichotomy similar to the concept of mitohormesis.^48–50^ Models in which ROS appear inhibitory, such as lung cancer and melanoma,^28,30,31^ may represent settings where cells already operate near the upper boundary of this window in contrast to cancer models showing a pro-migratory effect of mROS.^25,26^

Placing HIF-1a downstream of mROS and metabolic reprogramming in this signaling cascade has an important conceptual implication: metabolism does not merely provide fuel for a separately regulated migratory program but instead actively regulates it. Our findings suggest that the cell’s migratory “hardware”, cytoskeletal regulators, adhesion molecules, and motility factors, is transcriptionally controlled by the metabolic state through mROS/HIF-1a. This highlights a normoxic, redox-driven mode of HIF-1a activation that links mitochondrial metabolism directly to cell behavior and further supports the critical signaling role of mitochondria.^13,16,51^ Importantly, elevated glucose uptake in eMP-mROS^hi^ cells persists even in HIF-1a-knockout cells, confirming that the metabolic shift is upstream of and independent from HIF-1a signaling. Our identification of HIF-1a as a key mediator of metabolically driven lymphocyte migration is supported by recent work in both normal and malignant lymphoid cells. In regulatory T cells, HIF-1a functions as a metabolic switch directing glucose away from mitochondria toward glycolysis to support their migratory capacity despite enhancing their immunosuppressive function.^52^ In CLL, HIF-1a has been shown to regulate the expression of chemokine receptors and cell adhesion molecules and inactivation of HIF-1a impairs chemotaxis and stromal adhesion reducing bone marrow and spleen infiltration in xenograft models.^53^ Our data extend these findings by placing HIF-1a downstream of glucose metabolism and demonstrating that the mROS/HIF-1a axis transcriptionally controls the migratory program in lymphoid cancers under normoxic conditions.

The metabolic reprogramming we describe in malignant lymphocytes stands in contrast to the glucose allocation program that supports migration in normal T cells. Recent work has established that mitochondrial metabolism is the dominant pathway sustaining CD8+ and CD4+ T cell motility and migration. Simula et al. demonstrated that the TCA cycle, rather than glycolysis, is the principal metabolic pathway driving normal T cell migration and motility, with glucose and glutamine oxidation in mitochondria providing both the ATP and mROS required for motility.^35^ Complementing these findings, Steinert et al. showed that mitochondrial complex III function is essential for CD8+ T cell proliferation, naive cell maintenance, and memory formation, underscoring the broad dependence of normal T cell biology on intact mitochondrial electron transport and redox signaling.^54^

Our findings reveal a fundamentally different glucose allocation in malignant lymphocytes similar to an enhanced Warburg phenotype.^1,55,56^ Enhanced migratory mROS^hi^ cells show decreased glucose-carbon contribution to the TCA cycle due to downregulated citrate synthase and several other TCA cycle enzymes despite increased glucose uptake. In malignant lymphocytes, this divergent glucose allocation leads to elevated mROS and activation of an HIF-1a-dependent migratory program. In essence, malignant lymphocytes must divert glucose away from mitochondrial oxidation to maintain mROS/HIF-1a signaling and the migratory phenotype. Genetic deletion of citrate synthase recapitulates the highly migratory phenotype with enhanced migration and greater hepatic infiltration without affecting primary tumor growth. While the uncoupling of metabolic requirements of dissemination and proliferation is consistent with previously reported data our findings highlight the divergent metabolic reprogramming in lymphoid cancer. Whereas several solid tumors such as breast cancer, melanoma, squamous cell cancer rely on TCA cycle flux to fuel metastasis, malignant lymphocytes restrict glucose/pyruvate entry into the TCA cycle generating the mROS/HIF-1a signal that drives dissemination.^19,25,57–62^

Recent work has shown the importance of metabolic reprogramming and cellular ATP levels supporting lymphocyte migration and motility.^63^ Some studies show the importance of glycolytic ATP^52,64,65^ while others show the importance of OXPHOS-derived ATP in lymphocyte migration.^35,66–69^ Our data show that glycolytic ATP production is dispensable for malignant lymphocyte migration which is consistent with recently published work showing the critical importance of OXPHOS-derived ATP for T cell migration.^35^ Pyruvate supplementation alone restores motility in glucose-free conditions, and this rescue depends on pyruvate reduction-driven mROS/HIF-1a activation in the absence of glycolytic ATP. While this stands in contrast to some cancer models requiring glycolytic ATP for cell motility and metastasis,^70–72^ a unifying energetic strategy underlying metastasis has not emerged as mitochondrial respiration has also been shown to fuel invasion and metastatic dissemination in other tumor contexts.^57,73–75^ That the direction of pyruvate flux, not simply its abundance, controls migration is demonstrated by MCT1 knockdown experiments along with pharmacological modulation of pyruvate flux, where adding excess pyruvate does not rescue migration, ruling out a branch point-independent effect of pyruvate and confirming that it is the partitioning decision, not the substrate level, that matters. The pyruvate branch point thus operates as a true metabolic checkpoint with pyruvate conversion into lactate permitting mROS/HIF-1a-driven migration and oxidation in the TCA cycle suppressing migration. This extends the established regulatory role of pyruvate flux in solid cancer tumorigenesis and stem cell maintenance^7–9,11,12^ to the context of lymphoid cancer dissemination.

Convergent genetic, pharmacological, and metabolomic evidence presented here defines pyruvate metabolism as a previously unrecognized checkpoint for lymphoid cancer dissemination. AZD3965, which has completed Phase I dose-escalation studies in patients with advanced cancers including lymphoma and established a recommended Phase II dose with manageable on-target toxicities,^47^ reduces hepatic dissemination in our xenograft models and inhibits migration of primary CLL cells. These observations point to a new therapeutic rationale for MCT1 inhibition in lymphoid cancers: suppressing disease dissemination through lactate transport blockade by rerouting pyruvate flux into the TCA cycle. Whether this anti-migratory effect can be combined with conventional chemoimmunotherapy or other metabolic interventions to achieve durable control of lymphoid cancer spread warrants prospective investigation.

### Limitations of the study

Our mechanistic studies were largely performed in cell lines and xenograft models in immunodeficient mice and thus do not capture effects of the human immune microenvironment on metabolic regulation of dissemination. The glucose-dependent nature of the mROS states suggest that microenvironmental factors as well as metabolic factors at the organismal levels such as timing around meals or specific dietary composition could modulate this checkpoint *in vivo* in ways that our current models may not fully recapitulate. While we validate key findings in primary patient samples across CTCL, ALL, and CLL, validation in future human studies will be essential given the limitations discussed. At a mechanistic level, the upstream signals that suppress expression of citrate synthase and other TCA cycle enzymes underlying the metabolic reprogramming and fluctuations in migration potential and the specific HIF-1 target genes executing the pro-migratory transcriptional program remain to be identified. Elucidating the regulatory architecture around the pyruvate checkpoint will be critical for translating these findings into strategies that limit disease progression in lymphoid cancers.

## RESOURCE AVAILABILITY

Further information and requests for resources and reagents should be directed to and will be fulfilled by the Lead Contact, Stefan Schieke. This study did not generate new unique reagents.

## ACKNOWLEDGMENTS

We thank the Small Molecule Screening Facility at UW-Madison for assistance with Seahorse analyses, the Small Animal Imaging Facility at UW-Madison for their assistance with PET-CT studies, the Flow Cytometry Laboratory at UW–Madison, and the Flow Cytometry & Cell Sorting Shared Resource at GU-Lombardi Comprehensive Cancer Center for their assistance with analysis and cell sorting. This work was supported by a VA Merit Clinical Science Research & Development Grant I01CX002240 (S.M.S.), University of Wisconsin ICTR Translational Basic & Clinical Pilot Grant (S.M.S.), University of Wisconsin Skin Disease Research Center Pilot Grant (S.M.S.), and NIH grant R01CA266354 (L.R.). We used BioRender.com to create all visual graphics.

## AUTHOR CONTRIBUTIONS

Conceptualization: H.K., S.R., and S.M.S. Design of experiments: H.K., S.R., S.J., M.F., J.F., D.T.Y., S.M.S. Acquisition and/or analysis of data: H.K., S.R., S.J., M.F., N.M.H., P.B., A.P., A.N., D.T.Y., L.R., J.F., and S.M.S. Writing of the manuscript: H.K. and S.M.S., all authors have reviewed and/or revised the manuscript. Supervision: S.M.S.

## DECLARATION OF INTERESTS

The authors declare no competing interests.

## EXPERIMENTAL MODEL AND SUBJECT DETAILS

### Cell culture

The human malignant T-cell lines, Hut78, HH, Jurkat, and human malignant B-cell and myeloid lines Daudi, Toledo, Su-DHL-6, and HL-60 were obtained from the American Tissue Culture Collection (ATCC, Manassas, VA). Human CTCL line MyLa was generous gift from Dr. Reinhard Dummer, Dr. Emmanuel Contassot, and Dr. Lars French (University of Zurich). HH, Jurkat, Myla, Daudi and Toledo were propagated in RPMI-1640 media (ATCC) and Hut78 was propagated in Iscove’s Modified Dulbecco’s Medium (IMDM; ATCC). The following compounds were added to the media: 10% heat-inactivated fetal bovine serum (ATCC), 100 U/mL penicillin, 0.1 mg/mL streptomycin, and 0.1 mg/mL Amphotericin B (Gibco, Gaithersburg, MD). Cells were kept in a humidified incubator at 37°C with a 5% CO_2_ atmosphere. For hypoxic conditions, cells were transferred to a tri-gas incubator set to 3% O₂, 5% CO₂, and 92% N₂ to mimic hypoxic conditions. Cells were pre-incubated under hypoxia for 24 hours prior to migration assays. Where indicated, cells were incubated with phenformin (1-10 uM), N-acetylcysteine (4 mM), mitoTEMPO (30 uM), rotenone (10-500 nM), antimycin A (25-400 nM), 2-DG (25 mM), pyruvate (2.5-10 mM) all from Sigma-Aldrich, St. Louis; MO, PX-478 (30 uM), Oxamate (50 uM), Dichloroacetate (DCA) (10 uM), AZD3965 (1 uM), and IOX2 (50 uM), are from Selleckchem, Houston, TX.

### Mouse xenograft studies

For xenograft experiments, 2.5×10⁵ of sorted cells or 4×10⁶ of unsorted cells were injected subcutaneously into the bilateral flanks of 5-7 week-old immunodeficient NOD scid gamma (NSG) mice (NOD.Cg-Prkdcscid Il2rgtm1Wjl/SzJ) purchased from The Jackson Laboratory (Bar Harbor, ME). These studies were conducted in accordance with protocols approved by the Institutional Animal Care and Use Committees (IACUC) at both the University of Wisconsin-Madison and MedStar Health Research Institute in Washington, D.C. Where indicated, mice received daily intraperitoneal injections of either mitoTEMPO (100 mg/kg body weight), AZD3965 (100 mg/kg body weight), or a vehicle control, while NAC was provided at 1 g/L in their drinking water. Tumor growth was monitored daily tumor volume was calculated as V = D × 2d x 0.5, where D is the longest and d is the shortest diameter. When primary tumors reached an approximate volume of 1200 mm³, mice were euthanized, and their tumors and organs were collected.

### Human peripheral blood samples

De-identified blood samples from patients with acute lymphoblastic leukemia (ALL), chronic lymphocytic leukemia (CLL), from healthy subjects, and patients with leukemic cutaneous T cell lymphoma (CTCL, Sezary syndrome) were obtained with approval from the Health Sciences Institutional Review Board at the University of Wisconsin, Madison, WI and Institutional Review Board at Stanford University for all CTCL samples. The characteristics of these samples, including patient age and sex, are summarized in Tables S1 and S2. Samples of healthy adult volunteers served as controls (9 females and 4 males; mean age ± SD 46 ± 26.7 years). Leukemic burden was assessed via flow cytometry where indicated. For CLL, the markers included CD19^+^, CD20^dim^, CD5^+^, CD23^+^, CD200^+^, and kappa or lambda light chain restriction. For ALL, low side scatter, CD^45dim^, cytoplasmic CD3^+^, TdT^+^, with variable CD4, CD8, CD1a, and CD5 expression for T-ALL blasts, and CD19^+^, TdT^+^, with variable CD22, CD20, and CD79a expression for B-ALL blasts, were used. For CTCL/Sezary syndrome, CD4^+^, CD26^neg^ were used to determine circulating disease burden. Peripheral blood mononuclear cells (PBMCs) were isolated from the blood samples using a Ficoll (GE, Pittsburgh, PA) gradient centrifugation as per the manufacturer’s protocol. CD3^+^ and CD19^+^ cells were further isolated using MicroBeads labelled CD3^+^ and CD19^+^ (Miltenyi Biotec (Bergisch Gladbach, Germany) according to the manufacturer’s instructions. Isolated PBMCs were treated as indicated in the experiments.

### Mitochondrial ROS and cell sorting

Mitochondrial ROS levels were measured with MitoSox Red (Thermo Fisher Scientific Inc.) according to manufacturer’s instruction. Briefly, cells were stained with MitoSOX Red in HBSS buffer (Thermo Fisher Scientific Inc.) for 10 minutes at 37°C, followed by washing with HBSS buffer. Samples were acquired on Attune NxT Flow Cytometer (Thermo Fisher Scientific Inc.) for subsequent analysis. Isolation of cells with varying levels of mROS was carried out using a FACS Aria (BD Biosciences). For mROS-based sorting out of cell culture, cells kept in standard culture media were stained with MitoSox Red (Thermo Fisher Scientific Inc.) or CellROX Green (Thermo Fisher Scientific Inc.) according to manufacturer’s instructions followed by the addition of a cell viability dye such as SYTOX Green (Thermo Fisher Scientific Inc.) or propidium iodide (Thermo Fisher Scientific Inc.). For sorting from xenograft tumors, the tumors were finely chopped into smaller pieces and enzymatically digested with human Tumor Dissociation Kit (Miltenyi Biotec) to obtain single-cell suspensions for subsequent MitoSOX Red staining. For sorting of cells with high (mROS^hi^) and low (mROS^lo^) levels of mROS, approximately 5-10% of cells with the highest and lowest fluorescence intensity, respectively, we sorted. All sorted cells were collected in HBSS buffer (Thermo Fisher Scientific Inc.) for subsequent studies and analyses. Flow cytometric data was analyzed using FlowJo software (Tree Star Inc., Ashland, OR).

## METHOD DETAILS

### CRISPR/Cas9-knockout cell lines

HIF-1a was knocked out in the Hut78 cell line using the Edit-R system (Dharmacon, Lafayette, CO) as per the manufacturer’s guidelines and reported previously.^5^ Briefly, pre-designed CRISPR RNAs (crRNAs) targeting human HIF-1a were co-transfected into Hut78 cells alongside Edit-R transactivating CRISPR RNA (tracrRNA) and an Edit-R SMARTCas9 plasmid containing an mKate2 fluorescence marker. Transfected cells were then sorted based on mKate2 expression to enrich for edited cells using a BD FACS Aria (Becton Dickinson, San Jose, CA). For citrate synthase knockouts, Jurkat and HH cells were transduced with predesigned Edit-R Lentiviral sgRNA targeting human citrate synthase (Horizon Discovery) as per manufacturers instructions. Transfected cells were isolated based on eGFP expression using a BD FACS Aria (Becton Dickinson, San Jose, CA). Effective knockout was confirmed by western blot analysis demonstrating loss of HIF-1a or citrate synthase expression.

### siRNA-knockdown of MPC1 and MCT1

Transfections of MPC1 and MCT1 siRNA were performed in Jurkat and HH cell lines using Accell siRNA Reagents (Horizon Discovery), following the manufacturer’s instructions. A non-silencing control siRNA served as a negative control. Briefly, cells were transfected with target or control siRNA at a final concentration of 50 nM and cultured for 72 hours before further analysis. Knockdown of MPC1 and MCT1 was confirmed via western blot analysis.

### Plasmids and transfection

The plasmids for prolyl hydroxylation-defective HA-HIF-1a (P402A/P564A-pcDNA3, Addgene plasmid #18955) and wildtype HA-HIF-1a (pcDNA3, Addgene plasmid #18949) were generously provided by William Kaelin. Transfection of cells was carried out using the TurboFect Transfection Reagent (ThermoFisher Scientific, Waltham, MA) as per the manufacturer’s instructions. All experiments involving transfected cells were conducted three days post-transfection.

### Transwell migration/invasion assays

Matrigel-coated transwell inserts were utilized to assess cell migration, following the manufacturer’s protocol. In brief, 1×10⁵ cells were seeded into the transwell inserts and incubated for 24 hours. Serum-free medium was added to the upper chamber, while complete medium containing serum was placed in the lower chamber. After incubation, the cells that migrated to the lower chamber were counted and the percentage of migrated cells was calculated relative to the control group.

### Quantification of hepatic lymphocyte infiltration

To quantify malignant T cell infiltration in mouse organ tissue using flow cytometry, tissues were dissociated into single-cell suspensions using the human Tumor Dissociation Kit (Miltenyi Biotec). The resulting suspension was passed through a 70 µm filter to remove debris, then stained with human CD45 PE antibody according to manufacturer’s instructions (BioLegend, San Diego, California) for subsequent flow cytometric analysis.

To quantify malignant T cell infiltration in mouse organ tissue using light microscopy, xenograft liver tissues from indicated groups were preserved in 4% paraformaldehyde, followed by dehydration and permeabilization using gradient ethanol and xylene. The samples were then embedded in paraffin, and 2-µm-thick sections were prepared. These sections were deparaffinized with xylene and rehydrated through a graded ethanol series. After staining with hematoxylin and eosin, the sections were dehydrated again with gradient ethanol and xylene, sealed, and imaged under a microscope for further observation.

Liver tissue slides, stained with hematoxylin and eosin, were scanned at 20× magnification using an Aperio Digital Pathology Slide Scanner (Leica Biosystems), achieving a resolution of 0.5 mm per pixel and an image thickness of 3.5 mm. Intratumoral regions were selected and carefully annotated for analysis. The amount of lymphocyte infiltration was quantified as the percentage of area of infiltrating lymphocyte islands versus the entire liver section using the Aperio ImageScope version 12.4.3 (Leica Biosystems).

### Flow cytometric analysis of HIF-1a and glucose uptake

For HIF-1a quantification, cells were initially stained with CellROX Green (Thermo Fisher Scientific Inc.) as per the manufacturer’s instructions, followed by fixation with 4% formaldehyde. The fixed cells were then permeabilized and stained with HIF-1a antibody (BioLegend, San Diego, California). Glucose uptake was assessed using the Glucose Uptake Cell-Based Assay Kit (BioVision, Milpitas, CA), also in accordance with the manufacturer’s protocol.

Samples were acquired on Attune NxT Flow Cytometer (Thermo Fisher Scientific Inc.) and data was analyzed using FlowJo software (Tree Star Inc., Ashland, OR).

### Proliferation and cell viability analysis

For cell proliferation analysis, 25×10^4^ number of T-cells were seeded in triplicate in 24-well plates and cell counts were taken every 24 hours for 4 days after treatment with phenformin or with vehicle. For cell viability analysis, at the end of the experiments, cells were stained with SYTOX Green Nucleic Acid Stain (ThermoFisher Scientific, Waltham, MA) for 15 minute before flow cytometric analysis.

### Immunoblot analysis

Cells were lysed using radioimmunoprecipitation assay (RIPA) buffer supplemented with a protease inhibitor cocktail (Complete-mini, EDTA-free; Roche Applied Science, Indianapolis, IN). Protein concentrations were determined using the Bio-Rad Protein Assay (Bio-Rad, Hercules, CA). Nuclear extracts were isolated employing the Nuclear/Cytosol Fractionation Kit from BioVision (Milpitas, CA), following the manufacturer’s instructions. Proteins were then subjected to standard SDS-PAGE and Western blot analysis using antibodies against H2A-Z, β-Actin, HA tag, citrate synthase, MCT1, and MPC1 obtained from Cell Signaling Technology (Boston, MA) and HIF-1a from Novus Biologicals. H2A-Z, GAPDH, and β-actin were utilized as a loading control for nuclear fractions and whole cell lysates, respectively.

### Oxygen consumption analysis

The Seahorse XF-96 Flux Analyzer (Seahorse Bioscience, Billerica, MA) was utilized to measure the oxygen consumption rate (OCR) and extracellular acidification rate (ECAR). Briefly, 2×10⁵ sorted mROS^lo^ and mROS^hi^ Jurkat cells were seeded per well in 100 uL of sodium bicarbonate-free Dulbecco’s Modified Eagle’s Medium (DMEM) using a V3-PET cell culture plate. An additional 80 uL of DMEM was added to each well prior to conducting the XF analysis. Oxygen consumption and extracellular acidification were measured under basal conditions, with OCR values expressed as a percentage of the baseline oxygen consumption.

### Enzymatic assays

Citrate, pyruvate, lactate levels, and citrate synthase activity were quantified using the respective Colorimetric Quantitation Kit (BioVision, Milpitas, CA) following the manufacturer’s instructions. An equal number of cells from each experimental group were used for these analyses.

### Metabolomics and [U-^13^C]glucose tracing

For unlabeled metabolomic analyses, intracellular metabolites were extracted and analyzed from sorted cells in HBSS buffer following previously established methods.^76^ For isotope tracing experiments, unlabeled glucose was replaced with [U-^13^C]glucose at 25 mM in RPMI media. Cells were maintained in labelled media for 24 hours before cell sorting and subsequent analysis. After sorting, the cells were quickly pelleted at 1500 rpm for 5 minutes and the media was completely removed. Cell were washed with ice cold phosphate-buffered saline. The metabolites for each sample were extracted twice with cold LC-MS grade extraction solvent (40:40:20 acetonitrile:methanol:water, v:v) and the supernatants from cell extracts were combined. The metabolite extracts were dried under nitrogen stream, resuspended in LC-MS grade water, and analyzed using a Thermo Q-Exactive mass spectrometer coupled to a Vanquish Horizon UHPLC. The metabolites were separated on a 2.1 × 100 mm, 1.7 μM Acquity UPLC BEH C18 Column (Waters). The solvents used were A: 97:3 water:methanol (v:v), 10 mM tributylamine, 9 mM acetate, pH 8.2 and B: 100% methanol. The gradient was: 0 min, 95% A; 2.5 min, 95% A; 17 min, 5% A; 21 min, 5% A; 21.5 min, 95% A. The data were collected on a full scan negative mode. The metabolites identified were based on exact m/z and retention times that were determined with chemical standards. Peak integration was performed using MAVEN1. Relative metabolite levels were normalized to cellular protein content.

### RNAseq analyses

Total RNA was extracted using mRNA extraction kit (QIAGEN RNA extraction kit) according to the manufacturer’s protocol. Library and sequencing were prepared and analyzed by Novogene. For RNA-seq analysis, mapping, alignment, and differential expression were analyzed on the Galaxy public server (usegalaxy.org).^77^ Briefly, raw reads were aligned to the human reference genome (UCSC hg38) using HISAT2. Transcripts/genes were assembled with StringTie. Differential expressions were analyzed with DESeq2. GSEA was performed locally by GSEA software (V4.0.3) (http://software.broadinstitute.org/gsea/index.jsp). For the GSEA analysis, molecular signatures databases h.all.v5.2.symbols.gmt, KEGG were used. RNA-sequencing data discussed in this publication have been deposited in the National Center for Biotechnology Information’s Gene Expression Omnibus and are accessible through GEO Series accession number PRJNA1431357.

### Positron Emission Tomography/Computed Tomography (PET/CT) imaging

All mice were fasted for 12 hours prior to intravenous injection of approximately 8 MBq of 2′-deoxy-2′-(^18^F)fluoro-D-glucose (FDG) 1 hour before imaging. Mice were warmed after injection and anesthetized with inhalation gas using 2% isoflurane mixed with 1L/min of pure oxygen. Mice were imaged with the Siemens Inveon Hybrid microPET/CT (Siemens Medical Solutions, Knoxville, TN) in the prone position. Forty-million counts per mouse were collected for the PET scan to obtain adequate signal-to-noise. PET data were histogrammed into one static frame and subsequently reconstructed using ordered-subset expectation maximization (OSEM) of three dimensions followed by the maximum a posteriori algorithm, and CT attenuation and scatter correction were applied based on the NEMA NU 4 image-quality parameters. All PET and CT images were co-registered. Image data were analyzed using the General Analysis tools provided in the Siemens Inveon Research Workplace (Siemens Medical Solutions, Knoxville, TN). Data were identically window/leveled and scaled according to each animal’s decay corrected injection activity.

## QUANTIFICATION AND STATISTICAL ANALYSIS

Data are presented as the mean ± SD of at least triplicate measurements representative of 2 - 4 biologically independent experiments unless stated otherwise. Comparisons between different treatments were performed using student’s *t*-test and ANOVA for multiparametric comparisons in GraphPad Prism versions 7.0 and 8.0. A *p* value of less than 0.05 was considered statistically significant (* p < 0.05, ** p < 0.01, *** p < 0.001, *** p < 0.0001).

## DATA AND CODE AVAILABILITY

This study did not generate any unique datasets or code.

## Notes

### Competing Interest Statement

The authors have declared no competing interest.

## REFERENCES

1. Hanahan, D., and Weinberg, R.A. (2011). Hallmarks of Cancer: The Next Generation. Cell 144, 646–674. 10.1016/j.cell.2011.02.013.

2. Faubert, B., Solmonson, A., and DeBerardinis, R.J. (2020). Metabolic reprogramming and cancer progression. Science 368, eaaw5473. 10.1126/science.aaw5473.

3. Mao, Y., Xia, Z., Xia, W., and Jiang, P. (2024). Metabolic reprogramming, sensing, and cancer therapy. Cell Reports 43, 115064. 10.1016/j.celrep.2024.115064.

4. Vander Heiden, M.G., and DeBerardinis, R.J. (2017). Understanding the Intersections between Metabolism and Cancer Biology. Cell 168, 657–669. 10.1016/j.cell.2016.12.039.

5. Khan, H., Anshu, A., Prasad, A., Roy, S., Jeffery, J., Kittipongdaja, W., Yang, D.T., and Schieke, S.M. (2019). Metabolic Rewiring in Response to Biguanides Is Mediated by mROS/HIF-1a in Malignant Lymphocytes. Cell Reports 29, 3009–3018.e4. 10.1016/j.celrep.2019.11.007.

6. Khan, H., and Schieke, S.M. (2020). How to starve cancer cells when nutrients are abundant. Molecular & Cellular Oncology 7, 1718475. 10.1080/23723556.2020.1718475.

7. Olson, K.A., Schell, J.C., and Rutter, J. (2016). Pyruvate and Metabolic Flexibility: Illuminating a Path Toward Selective Cancer Therapies. Trends in Biochemical Sciences 41, 219–230. 10.1016/j.tibs.2016.01.002.

8. Flores, A., Schell, J., Krall, A.S., Jelinek, D., Miranda, M., Grigorian, M., Braas, D., White, A.C., Zhou, J.L., Graham, N.A., et al. (2017). Lactate dehydrogenase activity drives hair follicle stem cell activation. Nat Cell Biol 19, 1017–1026. 10.1038/ncb3575.

9. Schell, J.C., Wisidagama, D.R., Bensard, C., Zhao, H., Wei, P., Tanner, J., Flores, A., Mohlman, J., Sorensen, L.K., Earl, C.S., et al. (2017). Control of intestinal stem cell function and proliferation by mitochondrial pyruvate metabolism. Nat Cell Biol 19, 1027–1036. 10.1038/ncb3593.

10. Patel, M.S., and Rideout, T.C. (2026). Regulation of pyruvate dehydrogenase complex: Dancing to different drums in cancer. Intl Journal of Cancer 158, 1464–1480. 10.1002/ijc.70189.

11. Schell, J.C., Olson, K.A., Jiang, L., Hawkins, A.J., Van Vranken, J.G., Xie, J., Egnatchik, R.A., Earl, E.G., DeBerardinis, R.J., and Rutter, J. (2014). A Role for the Mitochondrial Pyruvate Carrier as a Repressor of the Warburg Effect and Colon Cancer Cell Growth. Molecular Cell 56, 400–413. 10.1016/j.molcel.2014.09.026.

12. Bader, D.A., Hartig, S.M., Putluri, V., Foley, C., Hamilton, M.P., Smith, E.A., Saha, P.K., Panigrahi, A., Walker, C., Zong, L., et al. (2018). Mitochondrial pyruvate import is a metabolic vulnerability in androgen receptor-driven prostate cancer. Nat Metab 1, 70–85. 10.1038/s42255-018-0002-y.

13. Martínez-Reyes, I., and Chandel, N.S. (2020). Mitochondrial TCA cycle metabolites control physiology and disease. Nat Commun 11, 102. 10.1038/s41467-019-13668-3.

14. Cheung, J.C.T., Ng, L.W., Zhu, Z., Chen, B., Li, S., Xu, M., Ding, X., Pu, D., Hu, Y., Ren, Y., et al. (2025). A Citrate Synthase Splice Variant Rewires the TCA Cycle to Promote Colorectal Cancer Progression. Cancer Research 85, 4450–4468. 10.1158/0008-5472.CAN-24-2355.

15. Schatton, D., and Frezza, C. (2025). Fine-tuning tumor immunogenicity with mitochondrial complex I. Nat Cancer 6, 231–233. 10.1038/s43018-024-00874-2.

16. Chandel, N.S. (2015). Evolution of Mitochondria as Signaling Organelles. Cell Metabolism 22, 204–206. 10.1016/j.cmet.2015.05.013.

17. Nunnari, J., and Suomalainen, A. (2012). Mitochondria: In Sickness and in Health. Cell 148, 1145–1159. 10.1016/j.cell.2012.02.035.

18. Vyas, S., Zaganjor, E., and Haigis, M.C. (2016). Mitochondria and Cancer. Cell 166, 555–566. 10.1016/j.cell.2016.07.002.

19. Bezwada, D., Perelli, L., Lesner, N.P., Cai, L., Brooks, B., Wu, Z., Vu, H.S., Sondhi, V., Cassidy, D.L., Kasitinon, S., et al. (2024). Mitochondrial complex I promotes kidney cancer metastasis. Nature 633, 923–931. 10.1038/s41586-024-07812-3.

20. Holmström, K.M., and Finkel, T. (2014). Cellular mechanisms and physiological consequences of redox-dependent signalling. Nat Rev Mol Cell Biol 15, 411–421. 10.1038/nrm3801.

21. Sena, L.A., and Chandel, N.S. (2012). Physiological Roles of Mitochondrial Reactive Oxygen Species. Molecular Cell 48, 158–167. 10.1016/j.molcel.2012.09.025.

22. Sullivan, L.B., and Chandel, N.S. (2014). Mitochondrial reactive oxygen species and cancer. Cancer Metab 2, 17. 10.1186/2049-3002-2-17.

23. Aurora, A.B., Khivansara, V., Leach, A., Gill, J.G., Martin-Sandoval, M., Yang, C., Kasitinon, S.Y., Bezwada, D., Tasdogan, A., Gu, W., et al. (2022). Loss of glucose 6-phosphate dehydrogenase function increases oxidative stress and glutaminolysis in metastasizing melanoma cells. Proc. Natl. Acad. Sci. U.S.A. 119, e2120617119. 10.1073/pnas.2120617119.

24. Brandl, N., Seitz, R., Sendtner, N., Müller, M., and Gülow, K. (2025). Living on the Edge: ROS Homeostasis in Cancer Cells and Its Potential as a Therapeutic Target. Antioxidants 14, 1002. 10.3390/antiox14081002.

25. Porporato, P.E., Payen, V.L., Pérez-Escuredo, J., De Saedeleer, C.J., Danhier, P., Copetti, T., Dhup, S., Tardy, M., Vazeille, T., Bouzin, C., et al. (2014). A Mitochondrial Switch Promotes Tumor Metastasis. Cell Reports 8, 754–766. 10.1016/j.celrep.2014.06.043.

26. Cheung, E.C., DeNicola, G.M., Nixon, C., Blyth, K., Labuschagne, C.F., Tuveson, D.A., and Vousden, K.H. (2020). Dynamic ROS Control by TIGAR Regulates the Initiation and Progression of Pancreatic Cancer. Cancer Cell 37, 168–182.e4. 10.1016/j.ccell.2019.12.012.

27. Labuschagne, C.F., Cheung, E.C., Blagih, J., Domart, M.-C., and Vousden, K.H. (2019). Cell Clustering Promotes a Metabolic Switch that Supports Metastatic Colonization. Cell Metabolism 30, 720–734.e5. 10.1016/j.cmet.2019.07.014.

28. Sayin, V.I., Ibrahim, M.X., Larsson, E., Nilsson, J.A., Lindahl, P., and Bergo, M.O. (2014). Antioxidants Accelerate Lung Cancer Progression in Mice. Sci. Transl. Med. 6. 10.1126/scitranslmed.3007653.

29. Le Gal, K., Ibrahim, M.X., Wiel, C., Sayin, V.I., Akula, M.K., Karlsson, C., Dalin, M.G., Akyürek, L.M., Lindahl, P., Nilsson, J., et al. (2015). Antioxidants can increase melanoma metastasis in mice. Sci. Transl. Med. 7. 10.1126/scitranslmed.aad3740.

30. Piskounova, E., Agathocleous, M., Murphy, M.M., Hu, Z., Huddlestun, S.E., Zhao, Z., Leitch, A.M., Johnson, T.M., DeBerardinis, R.J., and Morrison, S.J. (2015). Oxidative stress inhibits distant metastasis by human melanoma cells. Nature 527, 186–191. 10.1038/nature15726.

31. Wiel, C., Le Gal, K., Ibrahim, M.X., Jahangir, C.A., Kashif, M., Yao, H., Ziegler, D.V., Xu, X., Ghosh, T., Mondal, T., et al. (2019). BACH1 Stabilization by Antioxidants Stimulates Lung Cancer Metastasis. Cell 178, 330–345.e22. 10.1016/j.cell.2019.06.005.

32. Longo, J., DeCamp, L.M., Oswald, B.M., Teis, R., Reyes-Oliveras, A., Dahabieh, M.S., Ellis, A.E., Vincent, M.P., Damico, H., Gallik, K.L., et al. (2024). Glucose-dependent glycosphingolipid biosynthesis fuels CD8^+^ T cell function and tumor control. Preprint at Immunology, 10.1101/2024.10.10.617261.

33. MacIver, N.J., Michalek, R.D., and Rathmell, J.C. (2013). Metabolic Regulation of T Lymphocytes. Annu. Rev. Immunol. 31, 259–283. 10.1146/annurev-immunol-032712-095956.

34. Buck, M.D., Sowell, R.T., Kaech, S.M., and Pearce, E.L. (2017). Metabolic Instruction of Immunity. Cell 169, 570–586. 10.1016/j.cell.2017.04.004.

35. Simula, L., Fumagalli, M., Vimeux, L., Rajnpreht, I., Icard, P., Birsen, G., An, D., Pendino, F., Rouault, A., Bercovici, N., et al. (2024). Mitochondrial metabolism sustains CD8+ T cell migration for an efficient infiltration into solid tumors. Nat Commun 15, 2203. 10.1038/s41467-024-46377-7.

36. Frisch, A.T., Wang, Y., Xie, B., Yang, A., Ford, B.R., Joshi, S., Kedziora, K.M., Peralta, R., Wilfahrt, D., Mullett, S.J., et al. (2025). Redirecting glucose flux during in vitro expansion generates epigenetically and metabolically superior T cells for cancer immunotherapy. Cell Metabolism 37, 870–885.e8. 10.1016/j.cmet.2024.12.007.

37. Watson, M.J., Vignali, P.D.A., Mullett, S.J., Overacre-Delgoffe, A.E., Peralta, R.M., Grebinoski, S., Menk, A.V., Rittenhouse, N.L., DePeaux, K., Whetstone, R.D., et al. (2021). Metabolic support of tumour-infiltrating regulatory T cells by lactic acid. Nature 591, 645–651. 10.1038/s41586-020-03045-2.

38. Bailis, W., Shyer, J.A., Zhao, J., Canaveras, J.C.G., Al Khazal, F.J., Qu, R., Steach, H.R., Bielecki, P., Khan, O., Jackson, R., et al. (2019). Distinct modes of mitochondrial metabolism uncouple T cell differentiation and function. Nature 571, 403–407. 10.1038/s41586-019-1311-3.

39. Pucino, V., Certo, M., Bulusu, V., Cucchi, D., Goldmann, K., Pontarini, E., Haas, R., Smith, J., Headland, S.E., Blighe, K., et al. (2019). Lactate Buildup at the Site of Chronic Inflammation Promotes Disease by Inducing CD4+ T Cell Metabolic Rewiring. Cell Metabolism 30, 1055–1074.e8. 10.1016/j.cmet.2019.10.004.

40. Franchina, D.G., Dostert, C., and Brenner, D. (2018). Reactive Oxygen Species: Involvement in T Cell Signaling and Metabolism. Trends in Immunology 39, 489–502. 10.1016/j.it.2018.01.005.

41. Sena, L.A., Li, S., Jairaman, A., Prakriya, M., Ezponda, T., Hildeman, D.A., Wang, C.-R., Schumacker, P.T., Licht, J.D., Perlman, H., et al. (2013). Mitochondria Are Required for Antigen-Specific T Cell Activation through Reactive Oxygen Species Signaling. Immunity 38, 225–236. 10.1016/j.immuni.2012.10.020.

42. Scharping, N.E., Menk, A.V., Moreci, R.S., Whetstone, R.D., Dadey, R.E., Watkins, S.C., Ferris, R.L., and Delgoffe, G.M. (2016). The Tumor Microenvironment Represses T Cell Mitochondrial Biogenesis to Drive Intratumoral T Cell Metabolic Insufficiency and Dysfunction. Immunity 45, 374–388. 10.1016/j.immuni.2016.07.009.

43. Schieke, S.M., Phillips, D., McCoy, J.P., Aponte, A.M., Shen, R.-F., Balaban, R.S., and Finkel, T. (2006). The Mammalian Target of Rapamycin (mTOR) Pathway Regulates Mitochondrial Oxygen Consumption and Oxidative Capacity. Journal of Biological Chemistry 281, 27643–27652. 10.1074/jbc.M603536200.

44. Schieke, S.M., Ma, M., Cao, L., McCoy, J.P., Liu, C., Hensel, N.F., Barrett, A.J., Boehm, M., and Finkel, T. (2008). Mitochondrial Metabolism Modulates Differentiation and Teratoma Formation Capacity in Mouse Embryonic Stem Cells. Journal of Biological Chemistry 283, 28506–28512. 10.1074/jbc.M802763200.

45. Baumhoer, D., Tzankov, A., Dirnhofer, S., Tornillo, L., and Terracciano, L.M. (2008). Patterns of liver infiltration in lymphoproliferative disease. Histopathology 53, 81–90. 10.1111/j.1365-2559.2008.03069.x.

46. Bunchorntavakul, C., and Reddy, K.R. (2019). Hepatic Manifestations of Lymphoproliferative Disorders. Clinics in Liver Disease 23, 293–308. 10.1016/j.cld.2018.12.010.

47. Halford, S., Veal, G.J., Wedge, S.R., Payne, G.S., Bacon, C.M., Sloan, P., Dragoni, I., Heinzmann, K., Potter, S., Salisbury, B.M., et al. (2023). A Phase I Dose-escalation Study of AZD3965, an Oral Monocarboxylate Transporter 1 Inhibitor, in Patients with Advanced Cancer. Clinical Cancer Research 29, 1429–1439. 10.1158/1078-0432.CCR-22-2263.

48. Yun, J., and Finkel, T. (2014). Mitohormesis. Cell Metabolism 19, 757–766. 10.1016/j.cmet.2014.01.011.

49. Mitohormesis, an Antiaging Paradigm (2018). In International Review of Cell and Molecular Biology (Elsevier), pp. 35–77. 10.1016/bs.ircmb.2018.05.002.

50. Cheng, Y.-W., Liu, J., and Finkel, T. (2023). Mitohormesis. Cell Metabolism 35, 1872–1886. 10.1016/j.cmet.2023.10.011.

51. Chakrabarty, R.P., and Chandel, N.S. (2021). Mitochondria as Signaling Organelles Control Mammalian Stem Cell Fate. Cell Stem Cell 28, 394–408. 10.1016/j.stem.2021.02.011.

52. Miska, J., Lee-Chang, C., Rashidi, A., Muroski, M.E., Chang, A.L., Lopez-Rosas, A., Zhang, P., Panek, W.K., Cordero, A., Han, Y., et al. (2019). HIF-1α Is a Metabolic Switch between Glycolytic-Driven Migration and Oxidative Phosphorylation-Driven Immunosuppression of Tregs in Glioblastoma. Cell Reports 27, 226–237.e4. 10.1016/j.celrep.2019.03.029.

53. Valsecchi, R., Coltella, N., Belloni, D., Ponente, M., Ten Hacken, E., Scielzo, C., Scarfò, L., Bertilaccio, M.T.S., Brambilla, P., Lenti, E., et al. (2016). HIF-1α regulates the interaction of chronic lymphocytic leukemia cells with the tumor microenvironment. Blood 127, 1987–1997. 10.1182/blood-2015-07-657056.

54. Steinert, E.M., Furtado Bruza, B., Danchine, V.D., Grant, R.A., Vasan, K., Kharel, A., Zhang, Y., Cui, W., Szibor, M., Weinberg, S.E., et al. (2025). Mitochondrial respiration is necessary for CD8+ T cell proliferation and cell fate. Nat Immunol 26, 1267–1274. 10.1038/s41590-025-02202-x.

55. Murphy, M.P., and O’Neill, L.A.J. (2020). How should we talk about metabolism? Nat Immunol 21, 713–715. 10.1038/s41590-020-0691-8.

56. Vander Heiden, M.G., Cantley, L.C., and Thompson, C.B. (2009). Understanding the Warburg Effect: The Metabolic Requirements of Cell Proliferation. Science 324, 1029–1033. 10.1126/science.1160809.

57. LeBleu, V.S., O’Connell, J.T., Gonzalez Herrera, K.N., Wikman, H., Pantel, K., Haigis, M.C., De Carvalho, F.M., Damascena, A., Domingos Chinen, L.T., Rocha, R.M., et al. (2014). PGC-1α mediates mitochondrial biogenesis and oxidative phosphorylation in cancer cells to promote metastasis. Nat Cell Biol 16, 992–1003. 10.1038/ncb3039.

58. Christen, S., Lorendeau, D., Schmieder, R., Broekaert, D., Metzger, K., Veys, K., Elia, I., Buescher, J.M., Orth, M.F., Davidson, S.M., et al. (2016). Breast Cancer-Derived Lung Metastases Show Increased Pyruvate Carboxylase-Dependent Anaplerosis. Cell Reports 17, 837–848. 10.1016/j.celrep.2016.09.042.

59. Cai, Z., Li, C.-F., Han, F., Liu, C., Zhang, A., Hsu, C.-C., Peng, D., Zhang, X., Jin, G., Rezaeian, A.-H., et al. (2020). Phosphorylation of PDHA by AMPK Drives TCA Cycle to Promote Cancer Metastasis. Molecular Cell 80, 263–278.e7. 10.1016/j.molcel.2020.09.018.

60. Delaunay, S., Pascual, G., Feng, B., Klann, K., Behm, M., Hotz-Wagenblatt, A., Richter, K., Zaoui, K., Herpel, E., Münch, C., et al. (2022). Mitochondrial RNA modifications shape metabolic plasticity in metastasis. Nature 607, 593–603. 10.1038/s41586-022-04898-5.

61. Bartman, C.R., Weilandt, D.R., Shen, Y., Lee, W.D., Han, Y., TeSlaa, T., Jankowski, C.S.R., Samarah, L., Park, N.R., Da Silva-Diz, V., et al. (2023). Slow TCA flux and ATP production in primary solid tumours but not metastases. Nature 614, 349–357. 10.1038/s41586-022-05661-6.

62. Young, C.M., Beziaud, L., Dessen, P., Madurga Alonso, A., Santamaria-Martínez, A., and Huelsken, J. (2023). Metabolic dependencies of metastasis-initiating cells in female breast cancer. Nat Commun 14, 7076. 10.1038/s41467-023-42748-8.

63. Vuononvirta, J., Marelli-Berg, F.M., and Poobalasingam, T. (2021). Metabolic regulation of T lymphocyte motility and migration. Molecular Aspects of Medicine 77, 100888. 10.1016/j.mam.2020.100888.

64. Campello, S., Lacalle, R.A., Bettella, M., Mañes, S., Scorrano, L., and Viola, A. (2006). Orchestration of lymphocyte chemotaxis by mitochondrial dynamics. The Journal of Experimental Medicine 203, 2879–2886. 10.1084/jem.20061877.

65. Simula, L., Pacella, I., Colamatteo, A., Procaccini, C., Cancila, V., Bordi, M., Tregnago, C., Corrado, M., Pigazzi, M., Barnaba, V., et al. (2018). Drp1 Controls Effective T Cell Immune-Surveillance by Regulating T Cell Migration, Proliferation, and cMyc-Dependent Metabolic Reprogramming. Cell Reports 25, 3059–3073.e10. 10.1016/j.celrep.2018.11.018.

66. Chan, O., Burke, J.D., Gao, D.F., and Fish, E.N. (2012). The Chemokine CCL5 Regulates Glucose Uptake and AMP Kinase Signaling in Activated T Cells to Facilitate Chemotaxis. Journal of Biological Chemistry 287, 29406–29416. 10.1074/jbc.M112.348946.

67. Haas, R., Smith, J., Rocher-Ros, V., Nadkarni, S., Montero-Melendez, T., D’Acquisto, F., Bland, E.J., Bombardieri, M., Pitzalis, C., Perretti, M., et al. (2015). Lactate Regulates Metabolic and Pro-inflammatory Circuits in Control of T Cell Migration and Effector Functions. PLoS Biol 13, e1002202. 10.1371/journal.pbio.1002202.

68. Kishore, M., Cheung, K.C.P., Fu, H., Bonacina, F., Wang, G., Coe, D., Ward, E.J., Colamatteo, A., Jangani, M., Baragetti, A., et al. (2017). Regulatory T Cell Migration Is Dependent on Glucokinase-Mediated Glycolysis. Immunity 47, 875–889.e10. 10.1016/j.immuni.2017.10.017.

69. Ledderose, C., Liu, K., Kondo, Y., Slubowski, C.J., Dertnig, T., Denicoló, S., Arbab, M., Hubner, J., Konrad, K., Fakhari, M., et al. (2018). Purinergic P2X4 receptors and mitochondrial ATP production regulate T cell migration. Journal of Clinical Investigation 128, 3583–3594. 10.1172/JCI120972.

70. Shiraishi, T., Verdone, J.E., Huang, J., Kahlert, U.D., Hernandez, J.R., Torga, G., Zarif, J.C., Epstein, T., Gatenby, R., McCartney, A., et al. (2015). Glycolysis is the primary bioenergetic pathway for cell motility and cytoskeletal remodeling in human prostate and breast cancer cells. Oncotarget 6, 130–143. 10.18632/oncotarget.2766.

71. Zanotelli, M.R., Zhang, J., and Reinhart-King, C.A. (2021). Mechanoresponsive metabolism in cancer cell migration and metastasis. Cell Metabolism 33, 1307–1321. 10.1016/j.cmet.2021.04.002.

72. Zhan, H., Pal, D.S., Borleis, J., Deng, Y., Long, Y., Janetopoulos, C., Huang, C.-H., and Devreotes, P.N. (2025). Self-organizing glycolytic waves tune cellular metabolic states and fuel cancer progression. Nat Commun 16, 5563. 10.1038/s41467-025-60596-6.

73. Caino, M.C., Ghosh, J.C., Chae, Y.C., Vaira, V., Rivadeneira, D.B., Faversani, A., Rampini, P., Kossenkov, A.V., Aird, K.M., Zhang, R., et al. (2015). PI3K therapy reprograms mitochondrial trafficking to fuel tumor cell invasion. Proc. Natl. Acad. Sci. U.S.A. 112, 8638–8643. 10.1073/pnas.1500722112.

74. Caino, M.C., Seo, J.H., Wang, Y., Rivadeneira, D.B., Gabrilovich, D.I., Kim, E.T., Weeraratna, A.T., Languino, L.R., and Altieri, D.C. (2017). Syntaphilin controls a mitochondrial rheostat for proliferation-motility decisions in cancer. Journal of Clinical Investigation 127, 3755–3769. 10.1172/JCI93172.

75. Davis, R.T., Blake, K., Ma, D., Gabra, M.B.I., Hernandez, G.A., Phung, A.T., Yang, Y., Maurer, D., Lefebvre, A.E.Y.T., Alshetaiwi, H., et al. (2020). Transcriptional diversity and bioenergetic shift in human breast cancer metastasis revealed by single-cell RNA sequencing. Nat Cell Biol 22, 310–320. 10.1038/s41556-020-0477-0.

76. Britt, E.C., Qing, X., Votava, J.A., Lika, J., Wagner, A., Shen, S., Arp, N.L., Khan, H., Schieke, S.M., Fletcher, C.D., et al. (2023). Activation induces shift in nutrient utilization that differentially impacts cell functions in human neutrophils. Preprint at Biochemistry, 10.1101/2023.09.25.559385.

77. Afgan, E., Baker, D., Batut, B., van den Beek, M., Bouvier, D., Čech, M., Chilton, J., Clements, D., Coraor, N., Grüning, B.A., et al. (2018). The Galaxy platform for accessible, reproducible and collaborative biomedical analyses: 2018 update. Nucleic Acids Research 46, W537–W544. 10.1093/nar/gky379.

