## Supplemental Figures for "The pyruvate branch point controls lymphoid cancer cell dissemination"

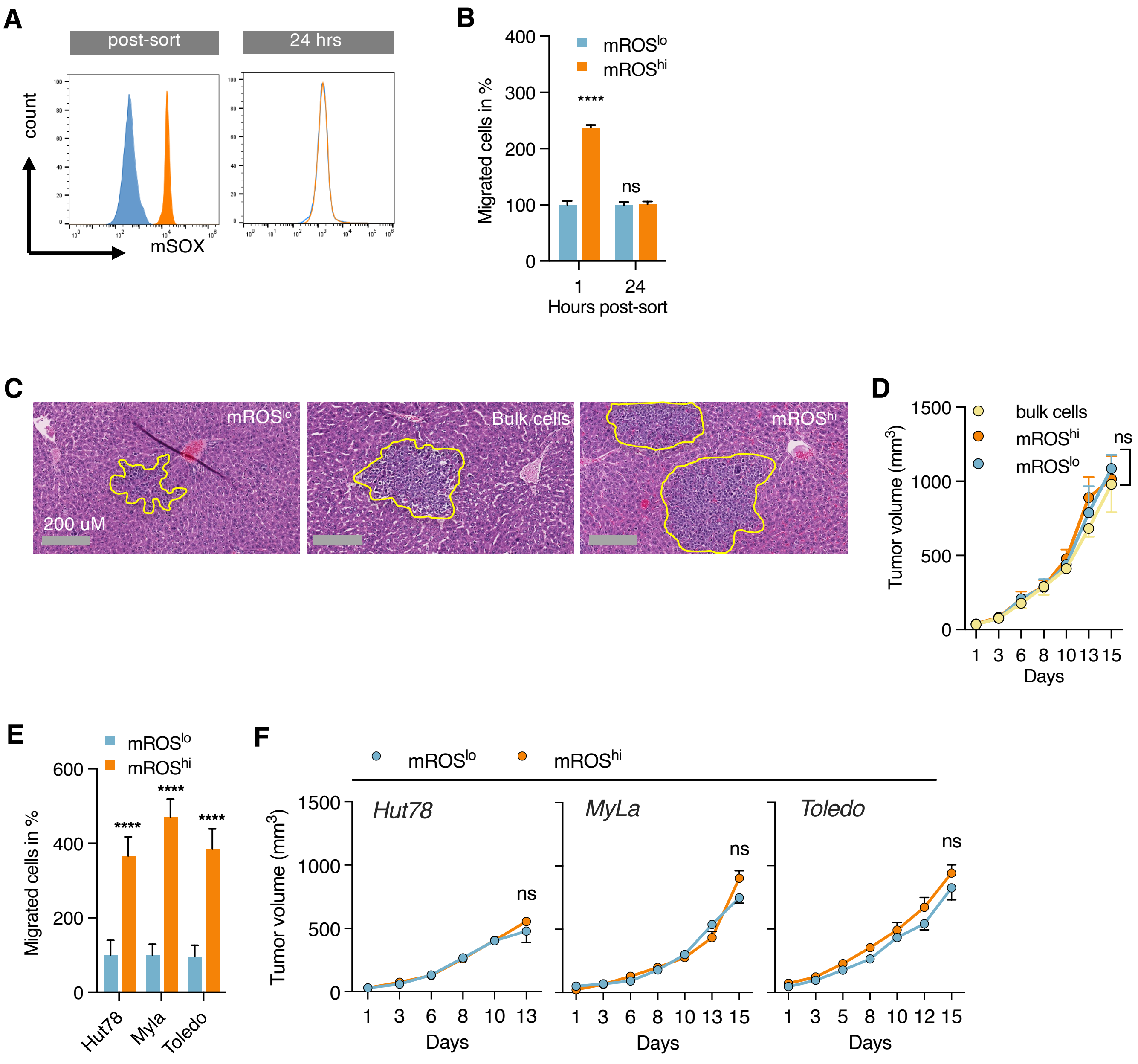


**Figure S1. Dynamic changes in mROS are linked to migration and hepatic infiltration but not primary xenograft tumor growth of malignant lymphocytes, related to Figure 1.** (A) mROS levels of isolated mROS$^{lo}$ and mROS$^{hi}$ Hut78 cells 1 hour after sorting and after 24 hours in culture. (B) Cell migration of mROS$^{lo/hi}$ Hut78 cells 1 hour and after 24 hours in culture. (C) Representative images of hematoxylin and eosin (H&E)-stained liver sections; infiltrating lymphocyte aggregates outlined in yellow. (D) Growth of xenograft tumors of sorted mROS$^{lo}$, mROS$^{bulk}$, and mROS$^{hi}$ Hut78 cells; n=4 tumors per group. (E) Transwell migration of isolated mROS$^{lo}$ and mROS$^{hi}$ xenograft tumor cells (n=6 tumor/group analyzed in triplicates). (F) Growth of second-generation xenograft tumors of sorted mROS$^{lo}$ and mROS$^{hi}$ cells (n=6 tumors). All data presented as mean ± SD of at least triplicate measurements.

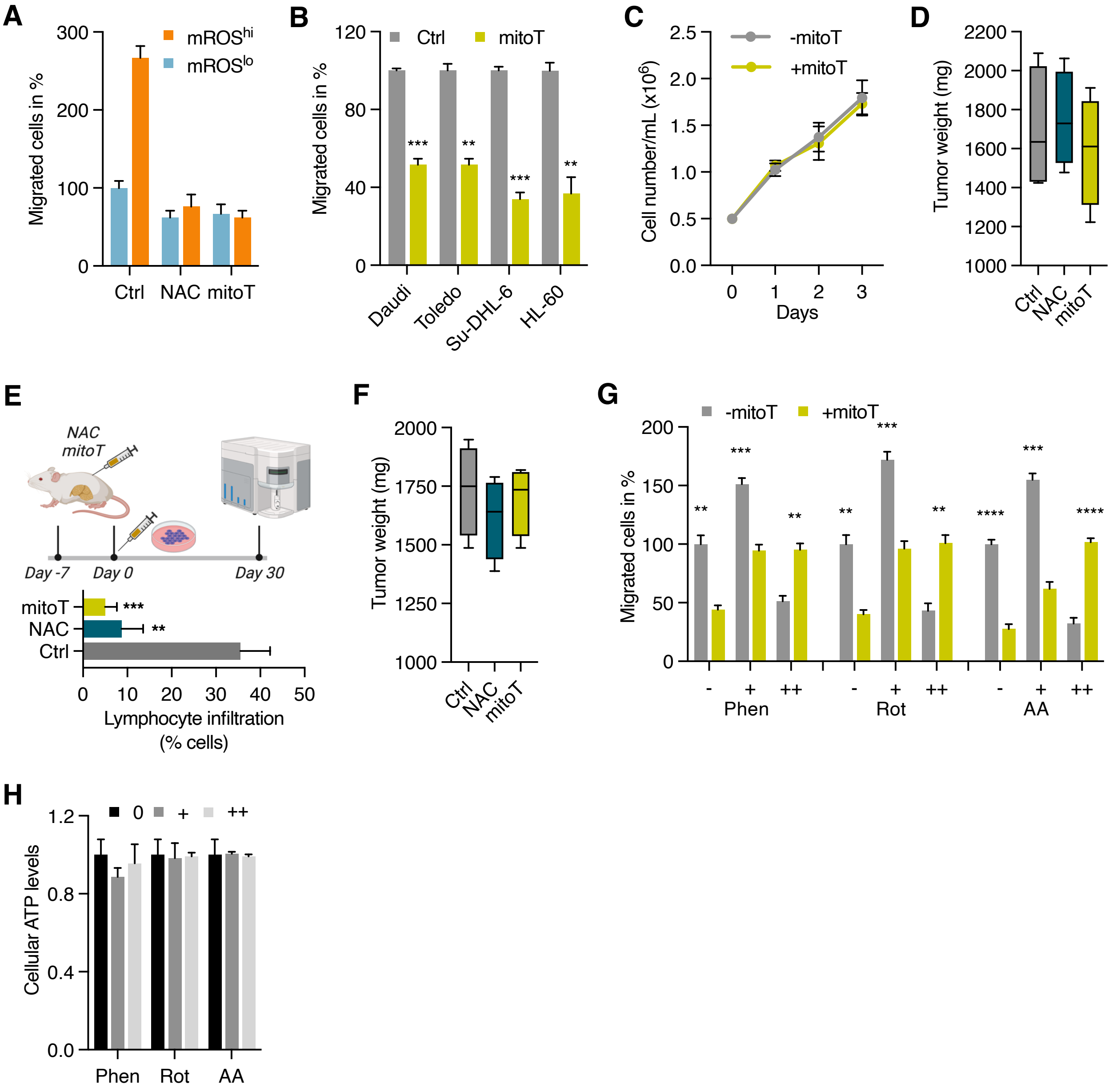


**Figure S2. Antioxidants and ETC inhibitors modulate migration and *in vivo* dissemination potential, related to Figure 1.** (A) Effect of antioxidants, N-acetylcysteine (NAC) and mitoTEMPO (mitoT), on transwell migration of sorted mROS$^{lo}$ and mROS$^{hi}$ Hut 78 cells based on CellROX Green fluorescence. (B) Effect of mitoT on transwell migration of indicated cell lines. (C) Effect of mitoT on doubling time of Hut 78 cells. (D) Weight of NAC and mitoTEMPO-pre-treated Hut78-derived xenograft tumors; n=4 tumors per group. (E) Hepatic dissemination of Hut78 cells in NAC- and mitoTEMPO-treated mice. Malignant lymphocytes in liver were quantified by flow cytometry using anti-hCD45 antibodies; n=3 mice/group. (F) Weight of NAC- and mitoTEMPO-pre-treated Hut78-derived xenograft tumors; n=4 tumors per group. (G) Effect of antioxidant mitoTEMPO on transwell migration of Hut78 cells treated with low (+) and high (++) concentrations of each phenformin (4 and 10 uM), rotenone (50 and 500 nM), antimycin A (100 and 500 nM). (H) Effect of low (+) and high (++) concentrations of phenformin, rotenone, and antimycin A on ATP levels in Jurkat cells (same concentrations as in G). All data presented as mean ± SD of at least triplicate measurements.

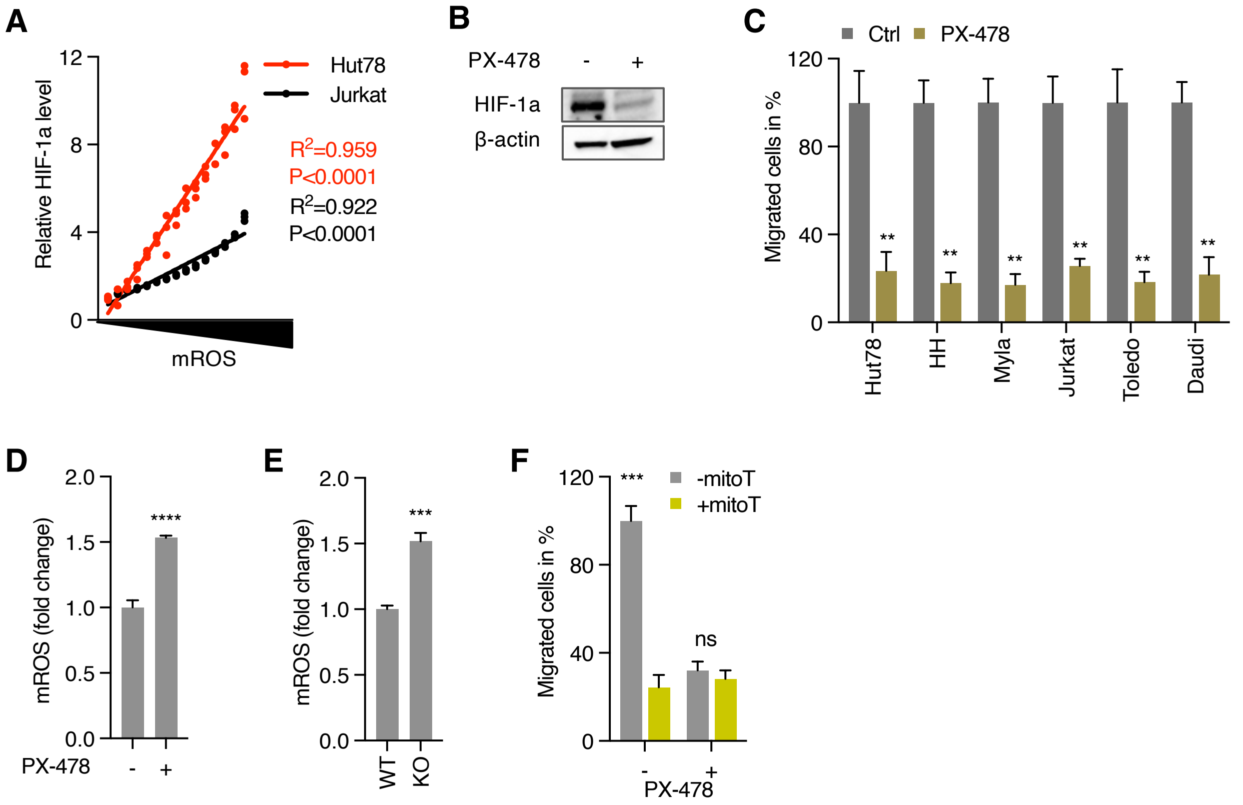


**Figure S3. Suppression of HIF-1a downstream of mROS reduces migration, related to Figure 2.** (A) mROS level and corresponding HIF1a expression in Hut78 and Jurkat cells quantified by flow cytometry. (B and C) Effect of HIF1a inhibition by PX-478 (B) on cellular migration in various cell lines (C). (D and E) Effect of HIF1a inhibitor, PX-478 (D) and HIF1a knockout (E) on mROS levels. (F) Effect of mitoTEMPO (mitoT) on transwell migration of Jurkat cells treated with PX-478. All data presented as mean ± SD of at least triplicate measurements.

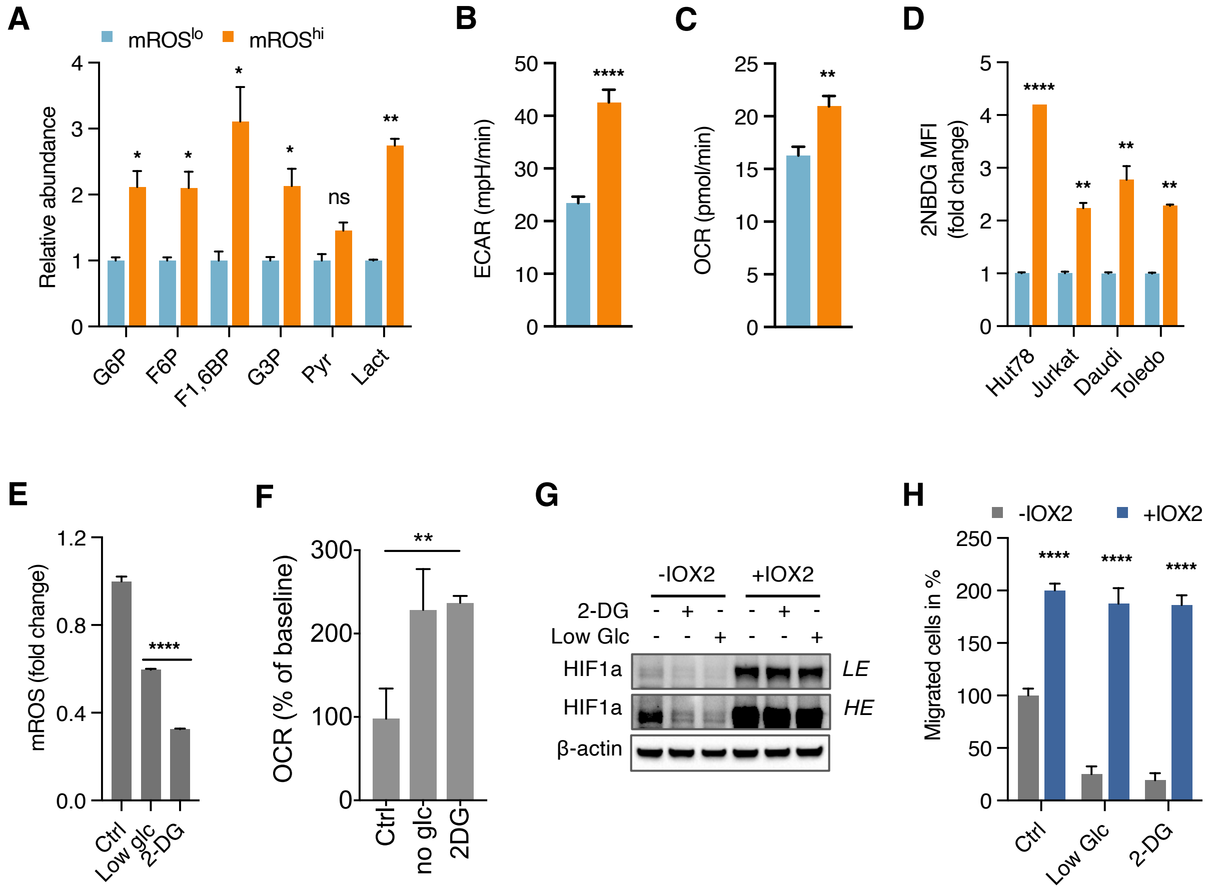


**Figure S4. Enhanced glucose uptake supports migration through HIF-1a, related to Figure 3.** (A) Relative abundance of glycolytic intermediate in metabolomic analyses of isolated mROS$^{lo}$ and mROS$^{hi}$ Jurkat cells. (B and C) Extracellular acidification rate (ECAR) and mitochondrial oxygen consumption rate (OCR) of isolated Jurkat cells. (D) Glucose uptake observed in isolated mROS$^{lo/hi}$ cells from indicated cell lines. (E) Effect of glucose-limiting conditions or inhibition of glycolysis with 2-deoxygluse (2-DG) on mROS levels. (F) Effect of glucose-limiting conditions or inhibition of glycolysis with 2-DG on oxygen consumption. (G and H) Immunoblots (G) and corresponding transwell migration assays (H) of Jurkat cells treated with PHD inhibitor, IOX2, in low glucose (3 mM) or 2-DG-containing media. mROS$^{lo}$ in light blue bars and mROS$^{hi}$ in orange bars in (A-D). All data presented as mean ± SD of at least triplicate measurements; ns, non-significant.

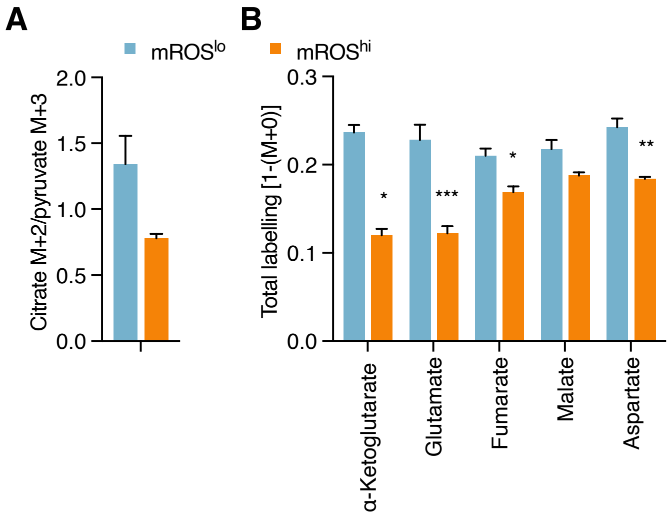


**Figure S5. Reduced glucose contribution to the TCA cycle in mROS$^{hi}$ cells, related to Figure 4.** (A) Ratio of [U-$^{13}$C]glucose-derived M+2 citrate and M+3 pyruvate. (B) Total labelling of TCA cycle metabolites [1-(M+0)] derived from [U-$^{13}$C]glucose. All data presented as mean ± SD of at least triplicate measurements.

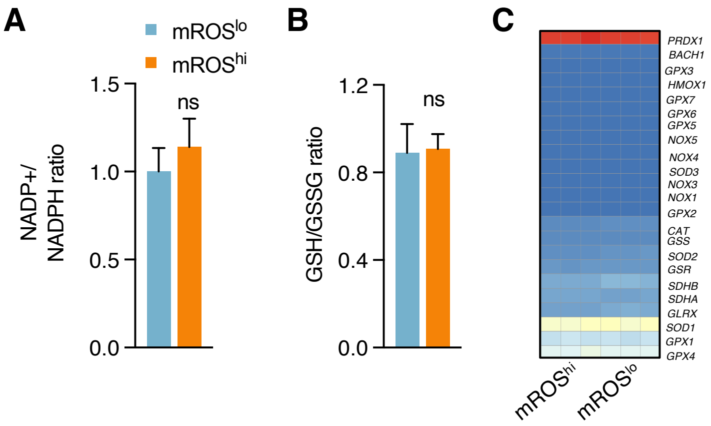


**Figure S6. Lack of activation of antioxidant defenses in mROS$^{hi}$ cells, related to Figure 4.** (A and B) Metabolic quantification of NADP+/NADPH ratio (A) and GSH/GSSG ratio (B) in sorted mROS$^{lo}$ and mROS$^{hi}$ Jurkat cells. (C) Clustered heatmap analysis of antioxidant gene profiles of isolated mROS$^{lo}$ and mROS$^{hi}$ Jurkat cells. All data presented as mean ± SD of at least triplicate measurements; ns, non-significant.

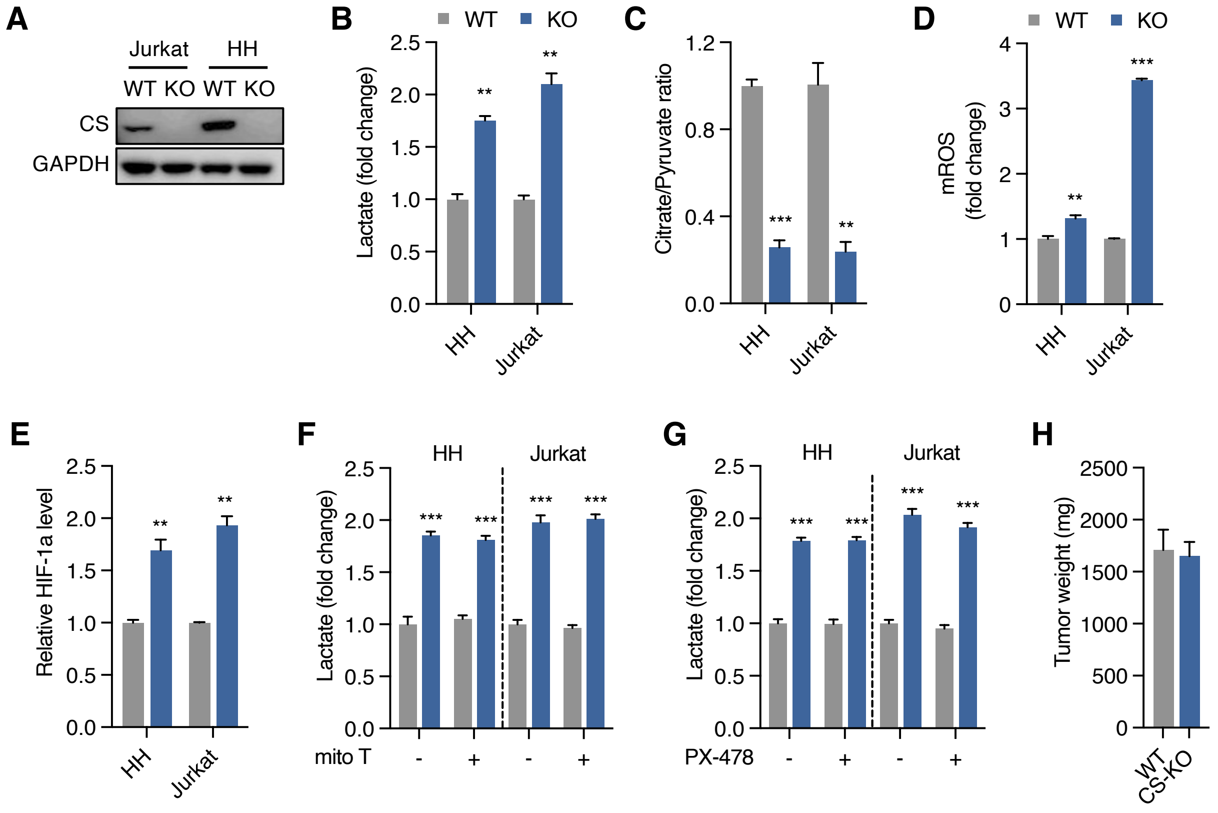


**Figure S7. Deletion of citrate synthase induces a glycolytic phenotype independent of mROS and HIF-1a, related to Figure 4.** (A) Immunoblots of CRISPR/Cas9-generated knockout (KO) clones show loss of citrate synthase (CS) expression. (B and C) Increased extracellular lactate level (B) and decreased citrate/pyruvate ratio (C) in CS-KO cell lines. (D and E) Increased mROS (D) and HIF-1a (E) levels in CS-KO cell lines. (F and G) Increased extracellular lactate levels are maintained in CS-KO cells in presence of mitoT (F) and PX-478 (G). (H) Weight of primary xenograft tumors derived from WT and CS-KO HH cells. Wildtype in grey bars and CS-KO in blue bars in (B-H). All data presented as mean ± SD of at least triplicate measurements.

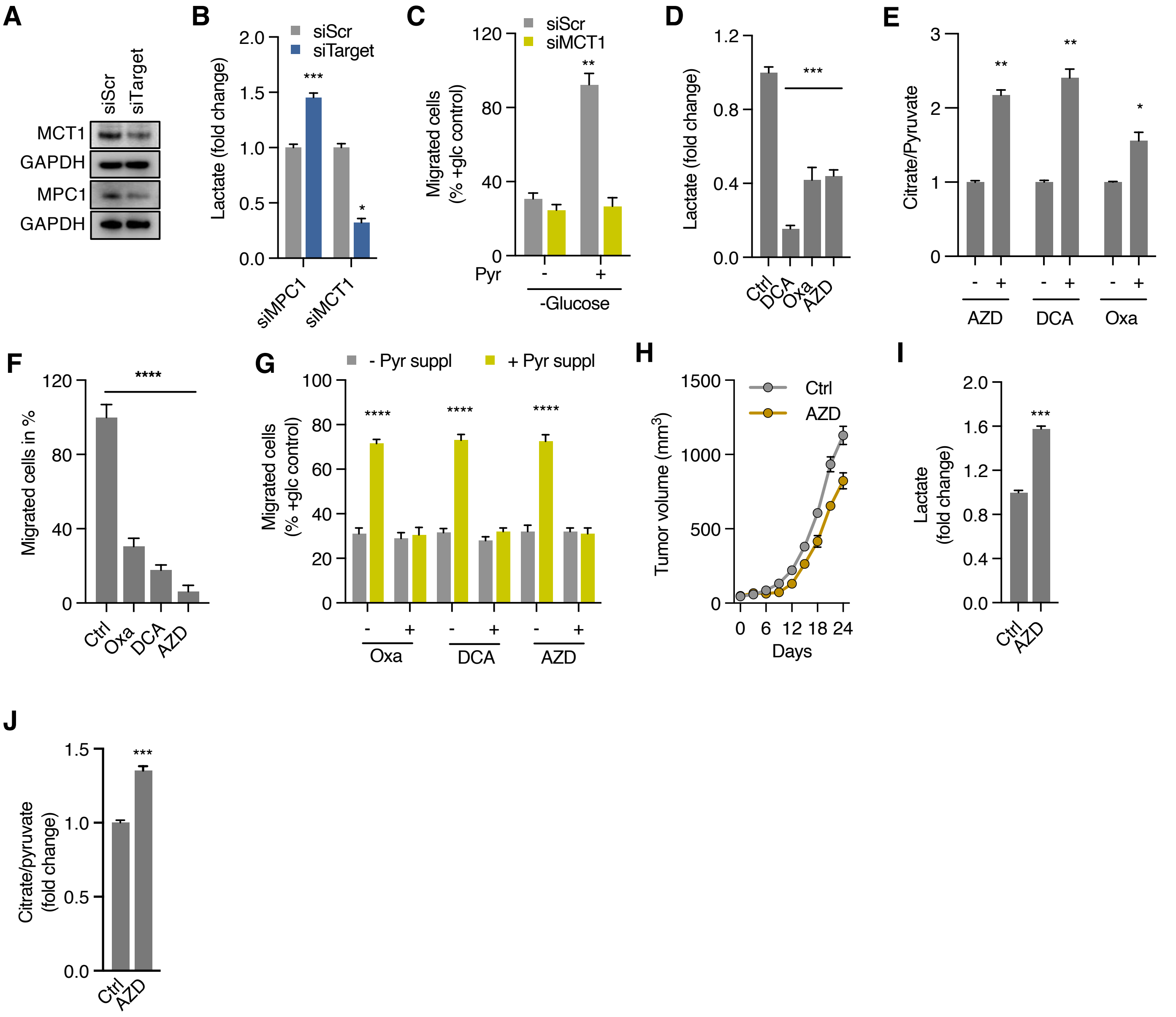


**Figure S8. Genetic and pharmacological modification of pyruvate flux regulates migration, related to Figure 5.** (A) Immunoblots show siRNA-mediated knockdown of MPC1 and MCT1. (B) Extracellular lactate levels of cells transfected with mitochondrial pyruvate carrier 1 siRNA (siMPC1), monocarboxylate transporter 1 siRNA (siMCT1) or scrambled siRNA (siSCR). (C) Effect of pyruvate supplementation on migration of cells transfected with monocarboxylate transporter 1 siRNA (siMCT1) or scrambled siRNA (siSCR) in glucose-free media. (D and E) Effect of DCA, oxamate, and AZD3965 on extracellular lactate levels (D) and citrate/pyruvate ratio (E). (F) Transwell migration of Jurkat cells treated with oxamate (Oxa), dichloroacetate (DCA), AZD3965 (AZD). (G) Effect of pyruvate supplementation on migration of Jurkat cells in glucose-free media in absence or presence of oxamate, dichloroacetate, and AZD3965. (I-K) Growth rate (I) of Hut-78-derived xenograft tumors in mice treated with AZD3965 versus control (n=4 tumors from 4 mice in each group) along with levels of lactate (J) and citrate/pyruvate ratio (K) in whole tumor lysates. All data presented as mean ± SD of at least triplicate measurements.

| Diagnosis | Age/Sex | WBC (1Kcells/uL) | Neutrophils | Lymphocytes | Monocyte | Eosinophils | Basophils | Leukemic burden |
|---|---|---|---|---|---|---|---|---|
| **B-Lymphoblastic Leukemia (ALL1)** | 18/M | 102.3 | 1% | 5% | 0% | 0% | 0% | 94% |
| **B-Lymphoblastic Leukemia (ALL2)** | 48/F | 10.4 | 8% | 12% | 1% | 1% | 0% | 77% |
| **B-Lymphoblastic Leukemia (ALL3)** | 63/F | 5.7 | 2% | 28% | 0% | 1% | 0% | 69% |
| **B-Lymphoblastic Leukemia (ALL4)*** | 10/F | - | - | - | - | - | - | 86% |
| **B-Lymphoblastic Leukemia (ALL5)** | 15/F | 3.7 | 10% | 25% | 1% | 0% | 0% | 63% |
| **B-Lymphoblastic Leukemia (ALL6)** | 65/F | 20.2 | 10% | 25% | 0% | 0% | 0% | 65% |
| **B-Lymphoblastic Leukemia (ALL7)** | 79/F | 9.1 | 6% | 27% | 0% | 0% | 0% | 67% |
| **T-Lymphoblastic Leukemia (ALL8)** | 71/M | 48.4 | 0% | 9% | 0% | 0% | 0% | 91% |
| **T-Lymphoblastic Leukemia (ALL9)** | 17/M | 933.9 | 1% | 3% | 1% | 0% | 0% | 95% |
| **T-Lymphoblastic Leukemia (ALL10)** | 30/F | 530.1 | 4% | 5% | 0% | 0% | 0% | 91% |
| **T-Lymphoblastic Leukemia (ALL11)** | 24/M | 103.3 | 13% | 5% | 4% | 0% | 0% | 76% |
| **Chronic Lymphocytic Leukemia (CLL1)** | 72/F | 82.4 | 12% | 87% | 1% | 0% | 0% | 79% |
| **Chronic Lymphocytic Leukemia (CLL2)** | 79/F | 26.5 | 12% | 86% | 2% | 0% | 0% | NA |
| **Chronic Lymphocytic Leukemia (CLL3)** | 64/M | 40.6 | 3% | 91% | 6% | 0% | 0% | 95% |
| **Chronic Lymphocytic Leukemia (CLL4)** | 49/M | 256.6 | 3% | 96% | 1% | 0% | 0% | 95% |
| **Chronic Lymphocytic Leukemia (CLL5)** | 62/F | 11.6 | 25% | 72% | 3% | 0% | 0% | 60% |
| **Chronic Lymphocytic Leukemia (CLL6)** | 64/M | 18.2 | 31% | 62% | 4% | 3% | 0% | 86% |
| **Chronic Lymphocytic Leukemia (CLL7)** | 72/M | 10.8 | 23% | 74% | 2% | 0% | 1% | 97% |
| **Chronic Lymphocytic Leukemia (CLL8)** | 58/M | 22.3 | 10% | 86% | 3% | 1% | 0% | 84% |
| **Chronic Lymphocytic Leukemia (CLL9)** | 46/M | 62.9 | 12% | 87% | 1% | 0% | 0% | 0% |
| **Chronic Lymphocytic Leukemia (CLL10)** | 72/M | 40.4 | 12% | 82% | 5% | 0% | 0% | 0% |
| **Chronic Lymphocytic Leukemia (CLL11)** | 90/M | 99.9 | 6% | 92% | 2% | 0% | 0% | 0% |
| **Chronic Lymphocytic Leukemia (CLL12)** | 71/F | 66.1 | 7% | 81% | 10% | 0% | 1% | 0% |
| **Chronic Lymphocytic Leukemia (CLL13)** | 83/M | 121.1 | 3% | 95% | 2% | 0% | 0% | 0% |

| | | | | | | | | |
|---|---|---|---|---|---|---|---|---|
| **Chronic Lymphocytic Leukemia (CLL14)** | 59/F | 26.1 | 24% | 70% | 4% | 1% | 0% | 0% |
| **Chronic Lymphocytic Leukemia (CLL15)** | 61/F | 41.6 | 13% | 85% | 1% | 1% | 0% | 0% |
| **Chronic Lymphocytic Leukemia (CLL16)** | 61/M | 175.3 | 8% | 92% | 0% | 0% | 0% | 0% |
| **Chronic Lymphocytic Leukemia (CLL17)** | 62/M | 1221% | 65% | 7% | 6% | 1% | 0% | 0% |
| **Chronic Lymphocytic Leukemia (CLL18)** | 69/M | 32.8 | 5% | 94% | 1% | 0% | 0% | 0% |
| **Chronic Lymphocytic Leukemia (CLL19)** | 53/M | 265.1 | 1% | 96% | 3% | 0% | 0% | 0% |
| **Chronic Lymphocytic Leukemia (CLL20)** | 66/M | 108.3 | 1% | 99% | 0% | 0% | 0% | 0% |
| **Chronic Lymphocytic Leukemia (CLL21)** | 70/F | 94 | 9% | 88% | 3% | 0% | 0% | 0% |
| **Chronic Lymphocytic Leukemia (CLL22)** | 63/F | 56.7 | 5% | 93% | 1% | 1% | 0% | 0% |
| **Chronic Lymphocytic Leukemia (CLL23)** | 89/F | 118.9 | 2% | 98% | 0% | 0% | 0% | 0% |
| **Chronic Lymphocytic Leukemia (CLL24)** | 48/M | 34.6 | 10% | 90% | 0% | 0% | 0% | 0% |
| **Chronic Lymphocytic Leukemia (CLL25)** | 70/F | 65 | 5% | 94% | 1% | 0% | 0% | 0% |

**Supplemental Table S1. Characteristics of leukemic samples.**
All acute lymphoblastic leukemia (ALL) and chronic lymphocytic leukemia (CLL) samples were collected from different patients with age and sex as indicated. Leukemic burden indicated as blasts as % of white blood cells (WBC) for ALL and as leukemic cells as % of lymphocytes for CLL. For standard immunophenotypic definition see STAR methods. *Bone marrow sample.

| Diagnosis | WBC (1Kcells/uL) | Lymphocytes | CD4+/CD26- (% of lymphs) | CD4+/CD26- (1Kcells/uL) |
|---|---|---|---|---|
| **Sézary Patient (CTCL1)** | 21.4 | 75% | 83% | 13.3 |
| **Sézary Patient (CTCL2)** | 16.8 | 50.3% | 55% | 4.6 |
| **Sézary Patient (CTCL3)** | 26.6 | 84.4% | 82% | 18.4 |
| **Sézary Patient (CTCL3)** | 10.6 | 34% | 75% | 2.7 |

**Supplemental Table S2. Characteristics of Sézary samples.**
All Sézary syndrome samples were collected from different patients.
